# Compound hypoxia with heat or acidification stress induces synergistic and additive effects on coral physiology

**DOI:** 10.64898/2025.12.01.690742

**Authors:** Kelly W. Johnson, Rebecca van Oostveen, Rene M. van der Zande, Noelle M. Lucey, Verena Schoepf

**Affiliations:** Department of Freshwater and Marine Ecology, Institute for Biodiversity and Ecosystem Dynamics, The University of Amsterdam, Amsterdam, The Netherlands; High Meadows Environmental Institute, Princeton University, New Jersey, United States; Department of Marine Sciences, University of Puerto Rico Mayagüez, Puerto Rico

**Keywords:** Hypoxia, ocean deoxygenation, climate change, ocean acidification, heat stress, compound climate events, multiple stressors, stressor interactions, coral bleaching, coral ecophysiology

## Abstract

As climate change accelerates, coastal marine ecosystems are increasingly exposed to co-occurring stressors whose combined effects are nonlinear and difficult to predict. Deoxygenation is a rapidly intensifying yet underrecognized threat to coral reefs that interacts with heat and acidification to alter coral physiology and stress resilience. However, the effects of hypoxia-related compound events on corals are largely unknown, underscoring the need for multi-stressor studies. Here, we conducted two extended-exposure experiments (12–17 days) across the coral species *Porites furcata*, *Porites astreoides* and *Siderastrea siderea*, to disentangle the individual and combined effects of low dissolved oxygen (hypoxia) with either heat or acidification. We measured eight phenotypic traits related to growth, metabolism, and symbiosis health to test whether hypoxia imposes energetic constraints or other physiological stress that amplify the effects of heat or acidification. Standardized effect size analysis across 24 stressor–trait combinations revealed 13 additive, 10 synergistic, and only one antagonistic response. Hypoxia consistently suppressed dark respiration by 37–49% across species and altered photophysiology in the two *Porites* species, whereas acidification alone had minimal effects, particularly in *S. siderea*. Heat stress caused the most pronounced declines across nearly all traits, and when combined with hypoxia, it produced the highest number of synergistic interactions. In contrast, the combination of hypoxia and acidification largely resulted in additive responses, suggesting that independent physiological mechanisms underlie these effects. All corals showed strong metabolic depression under hypoxia which is likely beneficial as a short-term adaptive response but may impose energetic constraints in the long-term. These findings highlight deoxygenation as critical yet often overlooked drivers of coral reef vulnerability. More multi-stressor experiments across a range of species are urgently needed to improve predictions of reef resilience under future ocean conditions, where compound stress events are expected to become more frequent and severe.

## Introduction

Ocean deoxygenation - the decline in dissolved oxygen (DO) - is emerging due to factors such as warming-driven changes in oxygen solubility and stratification as well as nutrient inputs (Breitburg et al., 2018; Keeling et al., 2009). Already, dramatic effects of reduced oxygen such as the formation of oxygen minimum zones are occurring in marine ecosystems, primarily reported in temperate zones (Diaz & Rosenberg, 2008) but increasingly also in tropical regions (Stramma et al., 2010). Furthermore, impacts of hypoxic events exceed the effects of ocean warming and acidification in many marine taxa (Sampaio et al., 2021). This is particularly concerning since deoxygenation rarely occurs in isolation but typically co-occurs with additional stressors, such as higher temperatures or low pH (Woods et al., 2022), also in the form of compound events (Gruber et al., 2021). This has led to recent calls for integrating aquatic deoxygenation into multi-stressor assessments such as the Planetary Boundary Framework (Ferrer et al., 2025; Rockström et al., 2009). Despite its significant ecological impact, deoxygenation has received comparatively less research attention than warming and acidification, particularly in tropical ecosystems where environmental variability is high and benthic organisms are operating near physiological thresholds (Ferrer et al., 2025; Gedan et al., 2017; Hughes et al., 2020; Woods et al., 2022).

Tropical coral reefs are biodiversity hotspots of immense socio-economic importance, yet deoxygenation has only recently been recognized as an emerging threat to reefs since they have traditionally been considered well-oxygenated (Altieri et al., 2017; Hughes et al., 2020; Pezner et al., 2023). Coastal tropical environments are especially vulnerable to acute deoxygenation (i.e., hypoxia, nominally defined as <2 mg L⁻¹) because shallow, low-flow conditions amplify the effects of warming, eutrophication, and elevated biological oxygen demand (Hughes et al., 2020; Nelson & Altieri, 2019). On coral reefs, prolonged hypoxia episodes have the potential to cause mass mortality and shifts in community structure, leading to so-called dead zones (Altieri et al., 2017). Coral reefs are further recognized as climate tipping elements (Armstrong McKay et al., 2022) that are susceptible to threshold-crossing events that drive mass coral bleaching or mortality (Johnson et al., 2021; Lucey et al., 2024). Tropical dead zones are likely far more widespread than current records indicate (Altieri et al., 2017), and projections suggest that up to 31% of reef sites could experience chronic, severe hypoxia by 2100 under high-emission scenarios (Pezner et al., 2023).

Extensive oxygen loss under rapid climate change is expected to interact with other climate stressors, likely compounding the impacts of warming and acidification and heightening the risk of reef ecological collapse (Deutsch et al., 2024). Consistent with these projections, the total number of hypoxic observations is expected to rise across even moderate warming scenarios by 2100 (Pezner et al., 2023). Hypoxic events will furthermore be directly linked with occurrences of low pH due to, for example, respiratory CO_2_ production, periods of low wind and thermal stratification, and eutrophication (Gobler & Baumann, 2016), as reported during a ∼6-day hypoxic event in Bocas del Toro, Panama (Johnson, Scott, et al., 2021). Together, these outcomes emphasize the urgency of determining how hypoxia interacts with other drivers, whether through additive or synergistic effects, since combined effects can be nonlinear. This priority is further stressed by the fact that the independent effects of hypoxia on corals remain severely understudied, as most experimental work has been limited to only a few species and short exposure durations (hours to a few days).

At the organism level, reef-building corals depend critically on oxygen to fuel host respiration and sustain their obligate symbiosis with dinoflagellates (*Symbiodiniaceae*), which drive photosynthetic energy production. Although corals produce O_2_ and, thus, tolerate strong diel oxygen fluctuations within their tissues (incl. hypoxia at night) (e.g., Linsmayer et al., 2020), recent studies reveal many species are nevertheless highly vulnerable to sustained low DO (e.g., Haas et al., 2014; Zhang et al., 2023). Yet, hypoxia remains among the least studied climate-linked stressors for corals, with only a few investigations into its combined effects with warming (e.g., Alderdice, Perna, et al., 2022; Lucey et al., 2024; Parry, Klein, & Duarte, 2025) or acidification (Osinga et al., 2017; Wijgerde et al., 2014). Most of these studies also used very short exposure durations (typically less than 12 hours; but see Altieri et al., 2017, Jain et al., 2023) and included limited biological responses. It is therefore largely unknown whether hypoxia exacerbates physiological stress caused by heat or acidification, and how this differs across species and physiological traits.

Current mechanistic models suggest that hypoxia and heat can activate overlapping bleaching cascades by depressing respiration, inducing oxidative stress, and disrupting metabolic compatibility between host and algal symbionts, increasing the likelihood that these cascades progress toward bleaching endpoints (Helgoe et al., 2024). Heat further raises metabolic oxygen demand, sharpening oxygen deficits under hypoxia and amplifying downstream host immune activation that can lead to exocytosis, apoptosis, or in situ degradation of symbionts (see references cited in Helgoe et al., 2024). This raises the likeliness that combined heat and hypoxia could produce synergistic outcomes where the combined impact is greater than the sum of each stressor on its own (i.e., additive effects). In contrast, hypoxia and acidification may act through largely distinct physiological pathways, as a recent meta-analysis reported mostly additive responses to deoxygenation–acidification interactions for performance, reproduction and survival in non-coral marine taxa (Steckbauer et al., 2020). However, the direction and magnitude of these interactions remain largely untested on corals, especially under prolonged exposures, limiting the ability to predict coral performance under increasingly frequent hypoxia-related compound events (Deutsch et al., 2024; Pezner et al., 2023).

To address these knowledge gaps, we investigated whether hypoxia intensifies the negative effects of warming and acidification on coral health and physiology through additive or synergistic effects. We conducted two separate, extended-exposure experiments (12-17 days) on three coral species representing diverse morphologies, assessing key functional traits including calcification, photosynthesis, respiration, and symbiont health. First, we hypothesized that oxygen limitation would constrain energy production (i.e., metabolic depression) and/or induce other physiological stress, thereby reducing the capacity to maintain the symbiosis and stable physiological function. Then, to resolve the nature and magnitude of combined effects, we utilized standardized effect sizes, allowing explicit classification of any interactive outcomes (i.e., non-additive responses). We hypothesized that stressors acting through overlapping physiological pathways (e.g., hypoxia and heat stress) would elicit synergistic effects, whereas those acting through distinct pathways (e.g., hypoxia and acidification stress) would produce additive effects.

## Materials and Methods

Two complementary, fully factorial experiments were conducted in 2023 on three coral species from two Caribbean reef systems (Figure 1). Experiment 1 tested the individual and combined effects of hypoxia and acidification on *Porites furcata* and *Siderastrea siderea* and took place at the Caribbean Research and Management of Biodiversity (CARMABI) Research Station in Curaçao, southern Caribbean. Experiment 2 examined the individual and combined effects of hypoxia and heat in *Porites astreoides*, and was conducted at the Smithsonian Tropical Research Institute (STRI) Research Station in Bocas del Toro, Panama. In both experiments, corals were exposed to one of four treatments: (1) ambient control, (2) hypoxia, (3) a secondary stressor (heat or acidification), or (4) the combined stressors. Hypoxia is defined here as DO at or below 2 mg L⁻¹, which refers to Experiment 2 and 1 respectively, based on widely used experimental applications across marine taxa reported in literature (Diaz & Rosenberg, 2008; Vaquer-Sunyer & Duarte, 2011). This hypoxia threshold is not intended to represent a species-specific survival limit in this study as these are unknown for most coral species. Corals were maintained under controlled laboratory conditions for 12 or 17 days for Experiment 1 and 2, respectively, placing these exposures at the upper extreme of durations used in previous hypoxia experiments, which range from less than one hour to only three studies lasting more than 12 days (Kvitt et al., 2022; Lucey et al., 2024; Mallon et al., 2025).

**Figure 1.**
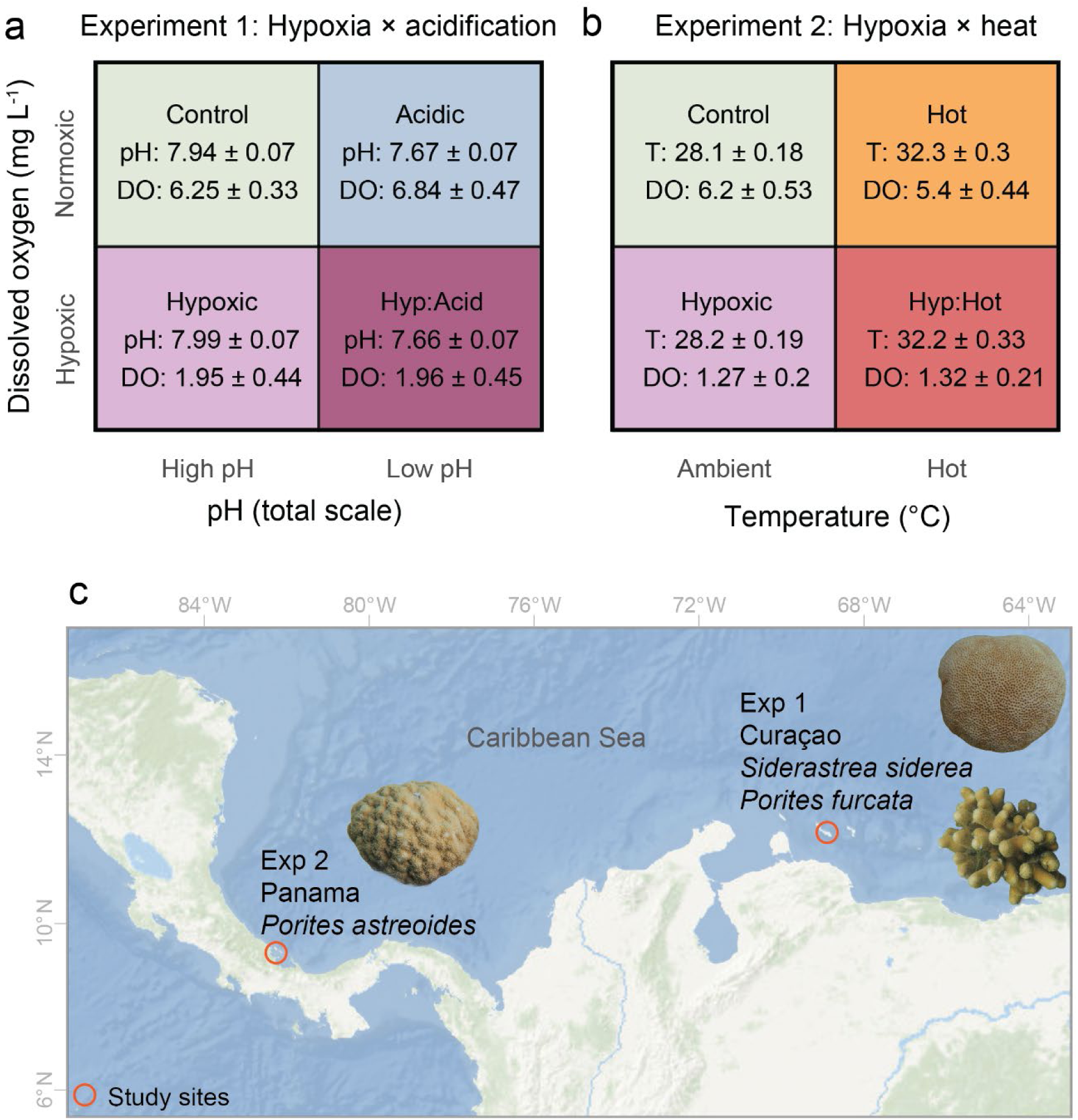
Experimental design and coral collection sites for two fully factorial stressor experiments. Abiotic conditions achieved in each experiment are shown, demonstrating the factorial combinations of stressors: (a) dissolved oxygen and pH in Experiment 1 (Southeastern coast, Curaçao), and (b) dissolved oxygen and temperature in Experiment 2 (Bocas del Toro, Panama). The map (c) shows the Caribbean with the two study locations highlighted, including inset images of the coral species used in each experiment: *Siderastrea siderea*, *Porites furcata*, and *Porites astreoides*.

### Experiment 1: Hypoxia × acidification

On 22 May 2023, six parent colonies of massive *S. siderea* and branching *P. furcata* were collected from Tugboat Reef (12°03′53.48″N, 68°51′36.69″W), Curaçao, under permit #2022/21467 issued to the CARMABI Foundation by the Curaçaoan Ministry of Health, Environment and Nature. Colonies were sampled from an average depth of 2.7–3.7 m for *S. siderea* and 3–5 m for *P. furcata*, with a minimum spacing of 5 m between colonies (genets) to maximize chances of collecting distinct genotypes.

The day after collection, each genet was fragmented into four ramets (one for each of the four treatments) and attached to acrylic tiles. These ramets were then returned to a site with environmental conditions similar to the collection location to recover under *in situ* conditions (4.7-m depth, CARMABI Reef; see Solomon et al., 2025). After a 10-day recovery period, they were transferred to the CARMABI Research Station and tank acclimated for 2 days in 26.3-L flow-through aquaria. Ramets from each genet were randomly assigned to one of four experimental treatments and distributed across replicate tanks, each consisting of different genet combinations, for a 17-20-day period starting on June 3: (1) Control, (2) Hypoxic, (3) Acidic, and (4) Combined Hypoxic and Acidic (Hyp:Acid) (Figure 1; SI Table 1), with two replicate tanks per treatment. Treatment target levels were 7.7 pH_T_ (acidic) and 2.0 mg O_2_ L^-1^ (hypoxic).

A complete list of treatment conditions and description of the setup are provided in SI Table 1 and SI Methods. In brief, corals were maintained in tanks (26.3-L) where temperature, DO, and pH_T_ were fully controlled and manipulated to generate the experimental stressor combinations (Apex Pro, Neptune Systems, USA). Temperature was continuously monitored and controlled by aquarium heaters, while DO and pH were regulated in real time using automated feedback systems that adjusted CO_2_, CO_2_-free air, and N_2_ gas injections. Tanks were supplied with 0.5 μm filtered seawater from a common reservoir, and daily discrete measurements were taken for temperature, pH_T_, DO, and salinity. Water samples for analysis of total alkalinity were collected periodically to document carbonate chemistry across tanks and treatments. Tank illumination followed a simulated natural day–night cycle (LED lights; maximum 376 ± 28 μmol photons m⁻² s⁻¹, *n* = 95; 13 h of daylight), and corals were fed live *Artemia* several times per week. Coral positions within tanks were rotated daily to minimize spatial variation in light exposure. This design allowed us to maintain precise and stable abiotic conditions while exposing corals to fully factorial combinations of DO and pH_T_, with stable temperature 27.6 °C.

### Experiment 2: Hypoxia × heat

Colonies of *Porites astreoides* exceeding 10 cm in diameter and exhibiting the brown color morph were collected on 4 October 2023, from Punta Caracol reef (9°22’40"N, 82°18’15"W; n = 8) under research and collection permit (Ministerio de Ambiente ARB-096-2023), and CITES export permits (Ministerio de Ambiente #PA01ARB-ARG219-2023). Colonies were sampled from an average depth of 3–4.5 m, with a minimum spacing of 10 m between colonies to reduce genotypic redundancy.

Each genet (n = 8) was fragmented into four ramets (n = 32), affixed to acrylic tiles (EcoTech Marine, USA), and allowed to recover for seven days under tank conditions: 30 °C and 100% air saturation (DO). Ramets were randomly assigned to one of four treatments for a 12-14-day period starting on 11 October 2023: (1) Control, (2) Hypoxic, (3) Hot, and (4) Combined Hypoxic and Hot (Hyp:Hot) (Figure 1; SI Table 1), with two replicate tanks per treatment. Treatment target levels were 32.5 °C (hot) and 1.3 mg O_2_ L^-1^ (hypoxic). Temperature in the Hot and Combined treatments was gradually increased from 30 °C to 32.5 °C by Day 7. Control and Hypoxic treatments were maintained at ∼28 °C. DO was gradually reduced to 2.5 mg L^-1^ on Day 1 and further decreased to the treatment target level of 1.3 mg L^-1^ on Day 2, which was then maintained thereafter.

Tank temperature and DO were controlled similarly to conditions in Experiment 1 to generate the experimental stressor combinations (Apex Pro, Neptune Systems, USA), and using the same automated feedback systems controlling temperature and DO (see SI Methods for more details). Although pH was not an experimental treatment in Experiment 2, it was continuously monitored and controlled using CO₂ gas addition to counteract increases from N₂ bubbling. Total alkalinity was measured periodically to report carbonate chemistry across tanks and treatments. Tanks were illuminated with a day–night cycle (LED lights; maximum 280 ± 29 μmol photons m⁻² s⁻¹, *n* = 114; 12 h of daylight), corals were fed live *Artemia* several times per week, and positions were rotated daily to minimize spatial variation in light exposure. This design maintained precise and stable abiotic conditions while exposing corals to fully factorial combinations of DO and temperature under controlled pH_T_.

#### Physiological analyses

To assess key functional traits associated with coral stress tolerance to hypoxia, heat, acidification, and their combinations, in both experiments we measured seven physiological metrics reflecting both coral host and algal symbiont (family Symbiodiniaceae) health and performance. Collectively, they provide a comprehensive understanding of productivity, energy metabolism, symbiosis integrity, and growth potential.

##### Photophysiology

*Symbiodiniaceae* photosynthetic performance was assessed by dark-adapted maximum quantum yield of photosystem II (F_v_/F_m_), measuring maximum potential for photochemical reactions using Pulse Amplitude Modulation (PAM) fluorometry. F_v_/F_m_ was repeatedly measured for all fragments in the same location on the fragment every two days following 30 minutes of dark acclimation (SI Figure 2; SI Methods for settings). For Experiment 1, measurements were taken at 19:00, except on day 8 when measurements failed due to technical error and were instead performed the following morning before the lights turned on. For Experiment 2, measurements were performed at 18:00. In both experiments, the optic cable was held at a fixed 1 cm distance from the coral surface, and the same location on each fragment was measured each time to ensure consistency. For *S. siderea* and *P. astreoides*, measurements were taken at the center of the colony; for *P. furcata*, the side of a branch was measured approximately 0.5–1 cm below the apical tip, favoring the top surface if the branch was angled.

##### Metabolic rates

Photosynthetic and respiratory rates of corals were assessed at the end of each experiment by incubating individual corals and measuring DO evolution/consumption. Prior to each measurement, corals were cleaned of algae and dark-acclimated for at least 30 minutes. They were then placed in acrylic incubation chambers (∼550–1030 mL depending on the height of the ramet) equipped with oxygen optodes connected via fiber optic cables to one of two multi-channel oxygen meter (OXY-4 SMA G3, PreSens GmbH, Germany), allowing for up to eight simultaneous measurements. Corals were allowed to acclimate in the chambers for at least 10 minutes prior to data collection.

Chamber starting seawater conditions matched those of the corals’ respective treatment tanks, with consistent light levels, temperature, pH, and initial DO concentrations. To maintain stable temperatures, chambers were partially submerged in a water bath fitted with a heater and circulation pump. Chambers were filled airtight with seawater from the respective treatment tanks, and each contained a magnetic stir bar to ensure thorough mixing for accurate oxygen measurements. Eight chambers were run concurrently, including one blank chamber containing only seawater as a background control. A 15-minute continuous segment of data was used to calculate oxygen evolution slopes.

In Experiment 1, oxygen content was recorded continuously during 30 minutes of dark respiration (R), followed by 40 minutes of light photosynthesis (P*_net_).* Oxygen rates were blank-corrected, standardized to seawater volume, and normalized to the coral surface areas (µmol min⁻¹ cm⁻²), from which gross photosynthesis (P_*gross*_) (1) and *P*: R ratios (2) were calculated as:

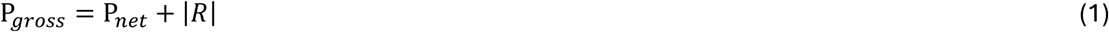

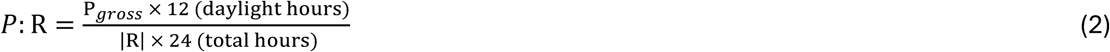

To account for each experimental coral, incubation trials/experiments began on Day 17 and continued over four consecutive days. Measurements proceeded by treatment group in the following order: Combined, Hypoxic, Acidic, and finally Control.

In Experiment 2, incubations were performed on a subset of genets (n = 6 of 8) across four consecutive days starting on Day 12. On the first two days, corals from the Hot and Combined treatments were measured, followed by Control and Hypoxic corals. Oxygen content was continuously recorded for 30 minutes of dark respiration (R), followed by 30 minutes of net photosynthesis (P*_net_),* and concluded with 10 minutes of light-enhanced dark respiration (LEDR), which was included only in Experiment 2. For Experiment 2, oxygen rates were blank-corrected, standardized to seawater volume, and normalized to the coral surface areas (µmol min⁻¹ cm⁻²), from which gross photosynthesis (*P*_*gross*_) (3) and P: R ratios (4) were calculated as:

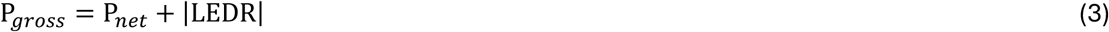

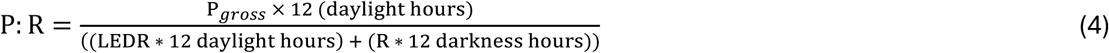

##### Calcification rates

Net calcification rates were measured from all coral fragments using the buoyant weight method (Jokiel et al., 1978) at two timepoints: (1) prior to the start of the experiment and (2) on the final day before oxygen incubations, to calculate weight change over the course of the experiment (Experiment 1: TA Plus 224 balance, Rice Lake Weighing Systems, USA; resolution: 0.0001 g; Experiment 2: Scout™ SPX222 balance, Ohaus Corporation, USA; precision: 0.01 g). Coral fragments were suspended by a monofilament line attached beneath the scale into a temperature-controlled seawater tank. Before weighing, all coral fragments were carefully cleaned of epiphytic algae.

Dry weights were calculated from buoyant weights using the formula from Jokiel et al. (1978), incorporating the densities of aragonite (2.93 g cm⁻³) and seawater (measured for each weighing session based on temperature and salinity) (HQ40d, HACH, USA). During the acclimation phase, high-resolution overlapping photographs of each coral fragment were taken from all directions at 0° and 45° angle relative to the fragment’s surface plane, and were processed into 3D models with AutoDesk ReCap Pro to measure live-tissue surface area (similar to Lavy et al., 2015). Changes in dry weight were normalized to live-tissue surface area and expressed as net calcification rates (mg day⁻¹ cm⁻²).

##### Symbiont density, total chlorophyll content, and tissue biomass

To assess symbiotic status and quantify coral bleaching, symbiont density and chlorophyll a + c2 content per algal symbiont cell (Symbiodiniaceae) were measured on frozen samples from the end of each experiment. Immediately following oxygen incubations, *S. siderea* and *P. furcata* fragments were stored at –80°C, while *P. astreoides* fragments were initially frozen at –20°C for three months before being transferred to –80°C for longer-term storage at the University of Amsterdam.

Coral tissue from a standardized 1 cm² area was removed from the skeleton using an airbrush (Master Performance S68 Multi-Purpose Precision Dual-Action Siphon Feed Airbrush, Master Airbrush, USA) followed by a water jet (Ultra WP-100, Waterpik, USA) with Milli-Q water (Milli-Q® Advantage A10, Merck KGaA, DE). The resulting tissue slurry was homogenized for 1 min (Tissue-Tearor, Biospec, OK, USA) and centrifuged at 4,000 × g for 10 minutes to isolate the symbiont pellet. The pellet was resuspended in 2 mL Milli-Q water and centrifuged again; the supernatant was discarded, and 5 mL Milli-Q water was added. This suspension was then partitioned for symbiont density counts (1 mL) and chlorophyll extraction (3 mL).

Symbiont densities were quantified using a Neubauer improved hemacytometer (Bright-Line™, Merck KGaA, DE) by counting six replicate 15 μL loadings under 100× magnification with a light microscope. The mean count per ramet was standardized to surface area.

For chlorophyll analysis, chlorophyll a and c₂ were extracted from the algal pellet in 100% acetone over 24 hours at 4°C in the dark. Absorbance was measured spectrophotometrically at 630, 663, and 770 nm to correct for turbidity (Novaspec Pro Spectrophotometer, Biochrom, MA, USA). Chlorophyll concentrations were calculated following Jeffrey & Humphrey (1975) and standardized to surface area (μg cm⁻²).

Tissue biomass was measured as ash-free dry weight (AFDW) of whole coral fragments with tissue, following a modified protocol from Fitt et al. (2000). Intact coral fragments (∼1 cm²), taken from branch tips or edges, were dried at 70°C until their weight stabilized, then combusted at 450°C for 6 hours to remove organic matter. Tissue biomass was standardized to surface area, determined using either caliper measurements or planar images analyzed with ImageJ (Schneider et al., 2012).

#### Data and statistical analyses

We used natural log response ratios (LnRR) to quantify the effects of individual and combined stressors because this metric enables intuitive interpretation of both directional and proportional change of biological variation relative to control conditions (Hedges et al., 1999). This was particularly important for identifying whether the effects of multiple stressors were additive, synergistic, or antagonistic (Crain et al., 2008; Steckbauer et al., 2020). The use of LnRR also facilitates direct comparisons across different species, response variables, and experimental settings, making it especially suitable for multi-species, multi-experiment studies conducted across geographic locations (Gurevitch et al., 2000; Nakagawa & Cuthill, 2007), as in the present study. Although linear mixed models can account for random effects and hierarchical structure, they were not the focus here, as our primary goal was to standardize and interpret effect sizes in a way that is broadly comparable and ecologically meaningful (Nakagawa & Schielzeth, 2013).

##### Standardizing and evaluating effect sizes across individual stressors

We quantified effect sizes of individual stressors using the natural log response ratio (LnRR), calculated as:

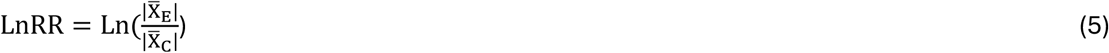

where the absolute value of X̄_E_ and X̄_C_ represent the mean response of the treatment and control groups, respectively. A positive LnRR indicates that the treatment increased the response variable relative to the control, while a negative value indicates a decrease. This metric enables consistent interpretation of treatment effects across varied biological endpoints.

To ensure precision across studies or replicates, each LnRR value was weighted by the inverse of its variance. The variance for each effect size was calculated as:

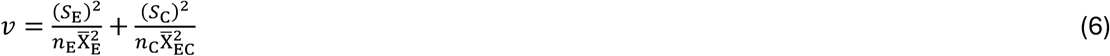

where *S*_E_ and S_C_ are the standard deviations, *n*_E_ and *n*_C_ are the sample sizes, and X̄_E_ and X̄_C_ are the means of the experimental and control groups, respectively.

Bias-corrected 95% confidence intervals (CIs) for each effect size were calculated using a bootstrap resampling procedure. CIs that did not overlap zero were considered statistically significant, indicating a positive or negative effect of the treatment relative to the control.

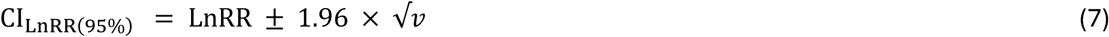

##### Evaluating combined stressor responses

To assess whether biological responses to combined stressors were additive or interactive (synergistic or antagonistic), we used a multiplicative null model. This approach compares the combined effect of two stressors against the expected additive effect of each stressor alone and is commonly applied in ecological meta-analyses, including marine invertebrates (Gurevitch et al., 2000; Harvey et al., 2013; Steckbauer et al., 2020).

Within this framework, the interaction effect size is calculated by comparing the log-transformed response of organisms exposed to both stressors to the sum of the individual stressor responses, while controlling for the response under baseline conditions. The equation used is:

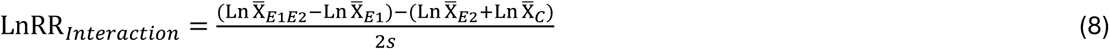

where X̄_*E*1*E*2_, X̄_*E*1_, X̄_*E*2_, and X̄_*C*_ represent the mean response under Combined, Hypoxic, secondary stressor (Acidic or Hot), and Control conditions, respectively.

Individual effects for each stressor were determined using the following equations:

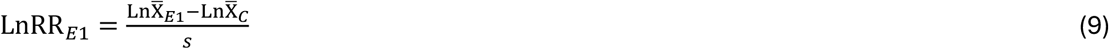

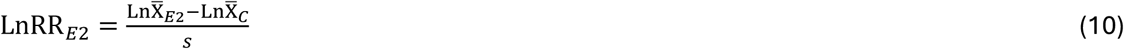

The pooled standard deviation *s* was calculated as:

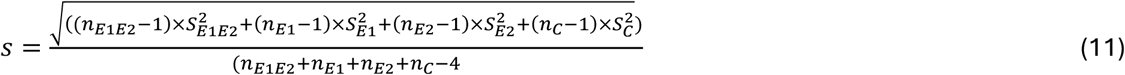

The variance associated with the interaction term was estimated using:

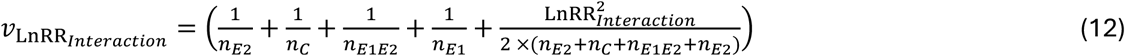

From this, 95% CIs were computed as:

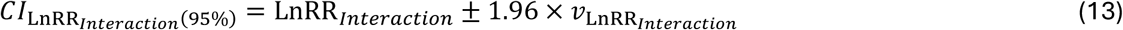

Interaction types were interpreted based on the 95% CI around the LnRR_*Interaction*_. If the *CI*_LnRR_*Interaction*(95%)__ overlapped zero, the response was considered additive, indicating acceptance of the null expectation that was calculated as the expected cumulative sum of individual stressor effects. Interactions were classified as synergistic when the combined effect exceeded the sum of the individual effects, and antagonistic when it fell below this expectation (Equation 8) (Gunderson et al., 2016; Todgham & Stillman, 2013)

The direction of the individual effects influence interpretation. When both individual effects were negative (or a mix of negative and positive), an interaction effect size less than zero indicated synergism, while a value greater than zero indicated antagonism. When both individual effects were positive, this logic was reversed: values greater than zero indicated synergism, and values less than zero were considered antagonistic, following the framework of Harvey et al. (2013). A conceptual graphic of these interaction types and their calculations is provided in SI Figure 1.

##### Predicted additive response

To assess the extent to which combined stressor responses deviated from additive expectations, we calculated a predicted additive response following the log-additive model (e.g. (Crain et al., 2008; Jackson et al., 2016). Specifically, the predicted response was computed as:

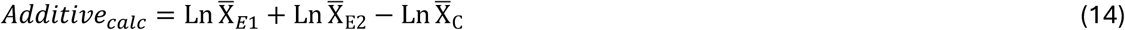

This additive prediction was then compared to the observed log-transformed response under the combined stressor treatment. The deviation between the observed and predicted values was calculated as follows:

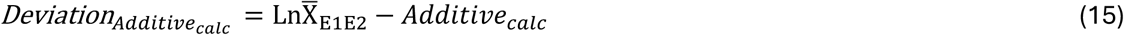

and the absolute magnitude of this deviation was used as a measure of how far the actual response departed from additive expectations.

## Results

### Individual stressor responses

Exposure to individual climate stressors (i.e., hypoxia, heat, or acidification) significantly affected coral physiology, but the direction and magnitude of responses varied by species and trait (Figure 2; SI Table 2). Analogous to mean ± error estimate, effect sizes are expressed here as LnRR ± precision, which is represented by half the width of the 95% confidence intervals. Further, percent changes are reported as the difference between treatment and control means, providing a quantitative, absolute measure of physiological response magnitude. Throughout both experiments, no tissue loss or mortality was observed in any coral ramets.

**Figure 2.**
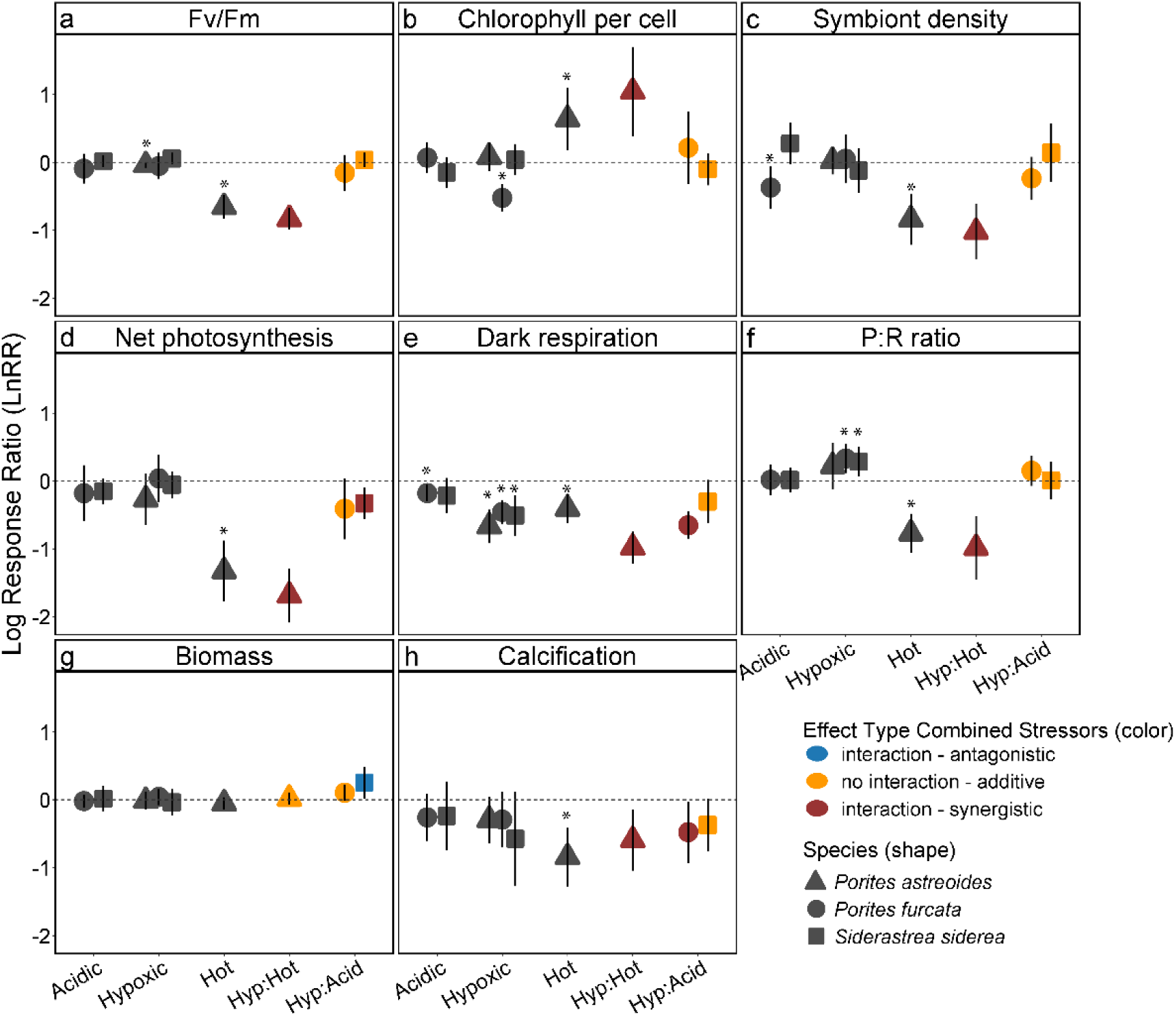
Log response ratios (LnRR, ± 95% confidence intervals (CI)) showing the effects of individual and combined stressors—hypoxia (hyp), heat, and acidification (acid)—on eight physiological traits across three Caribbean coral species. For individual stressors (grey symbols), a LnRR of zero indicates no effect, and positive or negative values indicate positive or negative responses, respectively. Symbol shapes represent coral species and asterisks indicate a significant effect of individual stressors (i.e., the CI does not overlap zero). For combined stressors (colored symbols), expected additive effects were calculated as the sum of individual LnRRs. Interaction effects represent deviations from these expectations: if the 95% confidence interval (CI) overlaps zero, the effect is additive (orange); otherwise, interactions are classified as synergistic (red) or antagonistic (blue) based on the direction of deviation. Traits analyzed include: (a) maximum quantum yield (Fv/Fm), (b) total chlorophyll content per symbiont, (c) symbiont density, (d) net photosynthesis (P), (e) dark respiration (R), (f) P:R ratio, (g) tissue biomass, and (h) net calcification.

#### Hypoxia

Hypoxia exposure broadly suppressed coral metabolic function, as indicated by negative LnRRs in all species where CIs did not overlap zero, although effect sizes differed across species (SI Table 2). Specifically, dark respiration declined by 37−49% under hypoxia in all three species (*P. astreoides*: 49% [–0.666 ± 0.243, LnRR ± half 95% CI], *P. furcata*: 37% [–0.456 ± 0.173], and *S. siderea*: 40% [–0.508 ± 0.297]). Suppressed respiration led to P:R ratios increasing by 40% in *P. furcata* (0.332 ± 0.214) and 33% in *S. siderea* (0.285 ± 0.217), but they were unaffected in *P. astreoides*, as indicated by a positive LnRR but with CIs overlapping zero (0.220 ± 0.342). Net photosynthesis rates and symbiont densities remaining unaffected across all three species (Figures 2–3).

**Figure 3.**
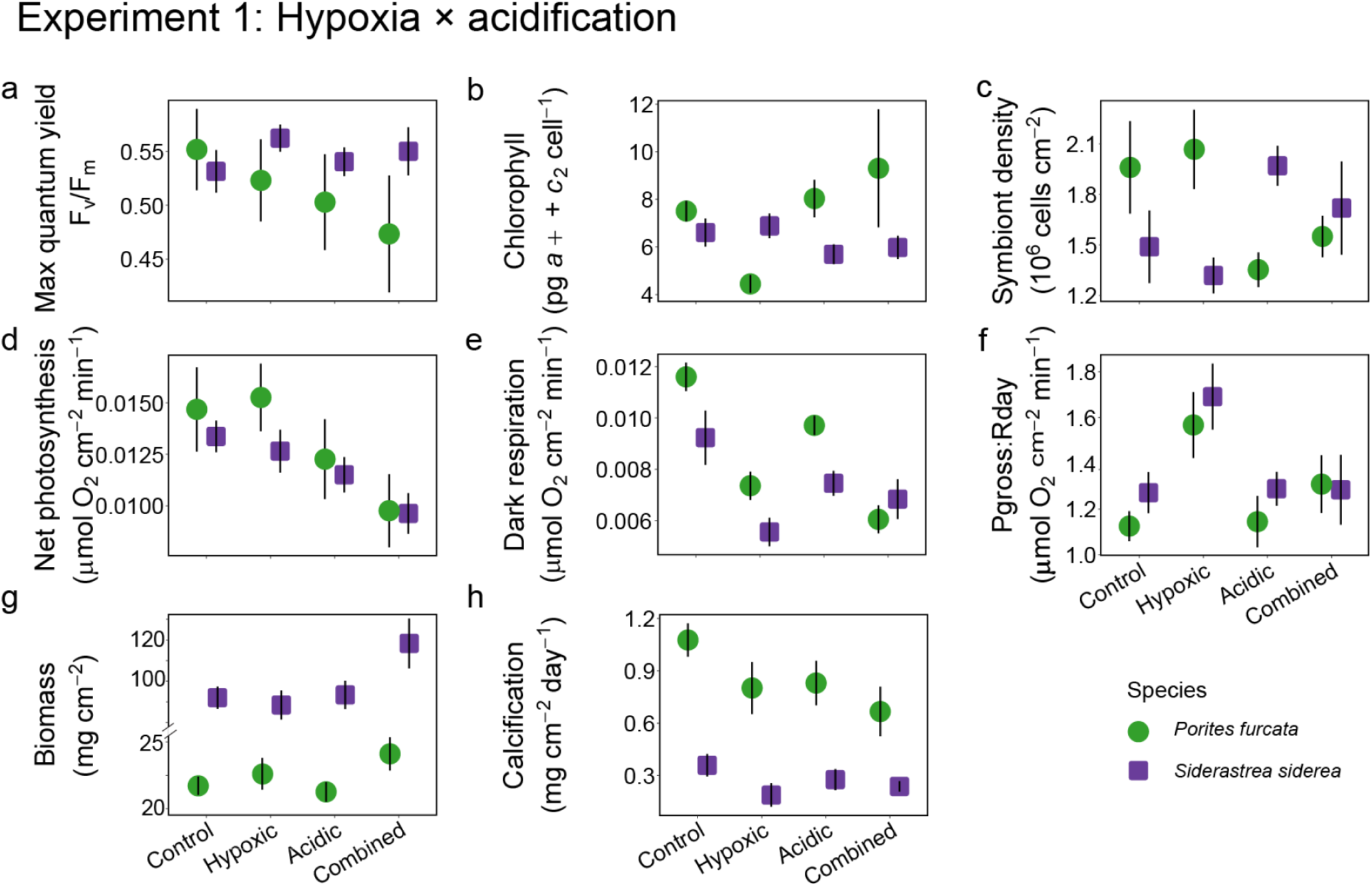
Experiment 1: Physiological responses of *Porites furcata* and *Siderastrea siderea* from Curaçao to the individual and combined effects of low dissolved oxygen (Hypoxic) and low pH (Acidic). Mean ± SE of raw values are shown. Corals were measured for endpoint performance metrics including a) maximum quantum yield (F_v_/F_m_), b) total chlorophyll content per symbiont cell, c) symbiont density per surface area, d) net photosynthetic rate, e) dark respiration rate (i.e., oxygen consumption), f) gross photosynthesis to daytime respiration ratio (P*_gross_*:R), g) tissue biomass and, h) net calcification.

Photosynthetic efficiency (F_v_/F_m_) declined by 4% in *P. astreoides* (–0.042 ± 0.041) (Figure 2, 4; SI Table 2) but remained stable in *P. furcata* (–0.053 ± 0.195) and *S. siderea* (0.056 ± 0.085) (Figure 2). Total chlorophyll content per symbiont cell declined by 41% in *P. furcata* (–0.524 ± 0.198) but remained unaffected for both *S. siderea* and *P. astreoides* (Figures 2–3). Combined with reduced respiration, this indicates a decoupling of autotrophic energy production and respiratory demand under oxygen stress. Net calcification and biomass, however, were not significantly affected by hypoxia in any species (Figure 2).

**Figure 4.**
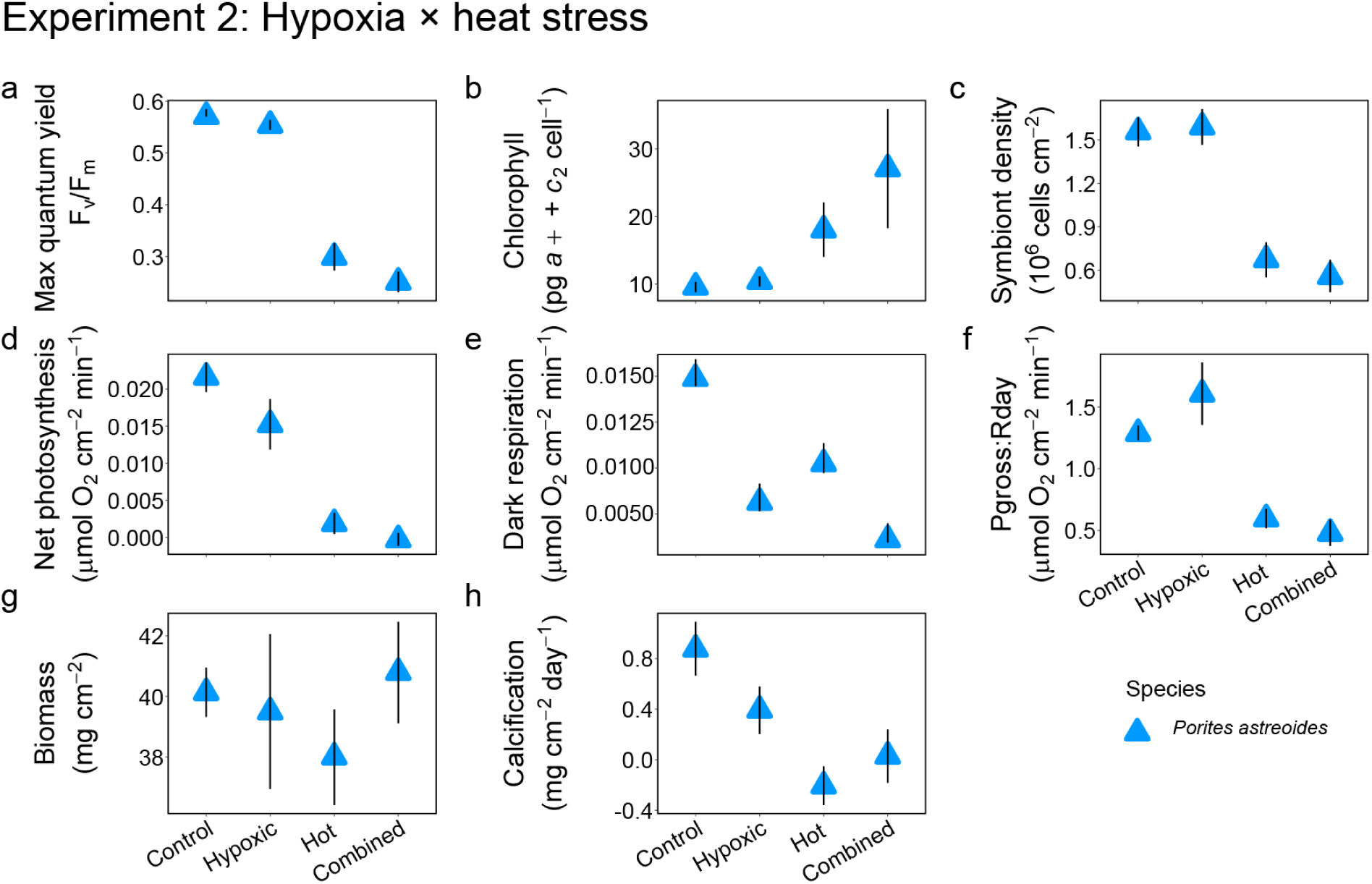
Experiment 2: Physiological responses of *Porites astreoides* from Panama to the individual and combined effects of low dissolved oxygen (Hypoxic) and high temperature (Hot). Mean ± SE of raw values are shown. Corals were measured for endpoint performance metrics including a) maximum quantum yield (F_v_/F_m_), b) total chlorophyll content per symbiont cell, c) symbiont density per surface area, d) net photosynthetic rate, e) dark respiration rate (i.e., oxygen consumption), f) gross photosynthesis to daytime respiration ratio (P*_gross_*:R), g) tissue biomass and, h) net calcification.

#### Acidification

Acidification alone had limited effects across species. Only *P. furcata* showed a 17% decline in dark respiration (–0.178 ± 0.120) and a 31% decline in symbiont density (–0.371 ± 0.312) (Figures 2–3). The 17% decline in respiration compares with a 37% decline under hypoxia, a 2.3-fold greater reduction (Figures 2–3). All other traits—including F_v_/F_m_, net photosynthesis, P:R ratio, calcification and biomass—remained unaffected in both *P. furcata* and *S. siderea* (Figure 2, SI Table 2). Notably, *S. siderea* exhibited complete physiological resistance to low pH, with all effect sizes being non-significant (i.e., CIs overlapping zero) across measured traits.

#### Heat

Heat stress caused widespread and severe physiological declines in *P. astreoides* (Figure 2, SI Table 2). All but one measured trait declined significantly, including calcification (45%; −0.845 ± 0.429), F_v_/F_m_ (57%; −0.657 ± 0.175), net photosynthesis (81%; −1.328 ± 0.448), P:R ratio (63%; −0.771 ± 0.289), dark respiration (63%; −0.405 ± 0.213), and symbiont density (64%; −0.840 ± 0.375) (Figure 2, 4). Biomass was the only trait that was unaffected (Figure 2, SI Table 2). Chlorophyll per cell was the only trait to increase substantially by 184% (Figure 4). Compared to hypoxia, heat stress reduced dark respiration by an added 13.9 percentage points and photochemical efficiency (F_v_/F_m_) by 52% more (Figure 4).

### Combined stressor responses

Interaction types varied by species and trait, revealing context-dependent stressor effects, with the magnitude of non-additive responses expressed as percent changes calculated from the difference between the observed combined treatment and the additive expectation of the individual stressor effects (Figures 3–4).

#### Additive responses dominate under hypoxia × acidification

Additive responses were most common under combined hypoxia and acidification in *P. furcata* and *S. siderea*, suggesting largely independent effects of each stressor. In *P. furcata*, six physiological traits exhibited additive effects: biomass (0.105 ± 0.118), chlorophyll per symbiont cell (0.215 ± 0.534), photochemical efficiency (F_v_/F_m_; –0.153 ± 0.261), P:R ratio (0.151 ± 0.221), net photosynthesis (–0.408 ± 0.448), and symbiont density (–0.235 ± 0.315) (Figure 2, SI Table 3).

*Siderastrea siderea* showed a similar pattern of additive effects across six parameters: calcification (–0.373 ± 0.392), dark respiration (–0.301 ± 0.316), F_v_/F_m_ (0.035 ± 0.108), chlorophyll per cell (–0.100 ± 0.237), symbiont density (0.144 ± 0.425), and P:R ratio (0.010 ± 0.271) (Figure 2, SI Table 3). One exception emerged for biomass in *S. siderea*, which showed an antagonistic interaction (0.251 ± 0.231), with the observed effect size being 6% smaller than predicted from additive expectations.

Despite the overall dominance of additive effects, some synergistic interactions were observed under hypoxia × acidification. In *P. furcata*, calcification (–0.479 ± 0.452) and respiration (–0.652 ± 0.200) declined 16% and 0.4% more than expected under the additive null, respectively, demonstrating that acidification - which triggered few significant responses in isolation - can exacerbate physiological stress when paired with hypoxia (Figure 2, SI Table 3). *Siderastrea siderea* exhibited a 3% decline in net photosynthesis (–0.328 ± 0.229), reflecting a measurable but moderate synergistic effect despite minimal response to acidification alone.

#### Synergistic physiological breakdown under hypoxia × heat

The combination of hypoxia and heat most often resulted in synergistic declines, with nearly all physiological traits measured in *P. astreoides* exhibiting deleterious effects under compounded stress beyond additive predictions (Figure 2, SI Table 3). Under compound stress, calcification synergistically declined 109% (−0.594 ± 0.452). Respiration likewise decreased synergistically 2% (–0.980 ± 0.234), exceeding the predicted additive effect by a modest but measurable amount of metabolic stress. Net photosynthesis, which had remained stable under hypoxia alone, was consistently suppressed synergistically by 2% under hypoxia and heat (–1.688 ± 0.396). This response was largely driven by the strong effect of heat stress alone (–1.328 ± 0.448) (SI Table 2). The reduction in photosynthesis under combined stress could be explained by the 54% decline in symbiont density (–1.023 ± 0.411), although chlorophyll per symbiont cell increased synergistically (11%, 1.044 ± 0.658), consistent with pigment retention or concentration within a reduced symbiont population. As a result of decreased net photosynthesis and respiration, the P:R ratio exhibited the most pronounced response with a synergistic decline (–0.985 ± 0.470) of 146%, underscoring the severity of compounded metabolic consequences. Photochemical efficiency also declined synergistically (11%, –0.833 ± 0.157). Finally, biomass (0.016 ± 0.090) exhibited an additive response, indicating that short-term growth was comparatively resistant to compound stress.

## Discussion

Hypoxia, warming, and acidification increasingly impact ocean systems globally both alone and in the context of compound stress events (Burger et al., 2022; Gruber et al., 2021). However experimental data addressing their combined effects on foundation species such as reef-building corals remain scarce, and the few available studies were typically very short in exposure duration, measured few traits, and often reported highly variable outcomes (e.g., Wijgerde et al., 2014; Osinga et al., 2017; Lucey et al., 2024). Here, by experimentally simulating hypoxia-related compound stress events lasting 12–17 days (Figure 1a–b), we demonstrate that coral responses are neither uniformly additive (the expected cumulative response of individual stressors) nor synergistic (where effects exceed the expected additive response). Instead, these responses span a spectrum of interactions that vary with stressors, species and traits (Figure 2, SI Table 2–3). Specifically, across the 24 combined stressor effect sizes measured in three coral species, 10 were synergistic, 13 were additive, and only one was antagonistic.

### Synergistic outcomes dominate when hypoxia is combined with heat

When hypoxia was combined with thermal stress, it elicited pronounced synergistic declines across most measured traits, including net photosynthesis, dark respiration, symbiont density, and calcification. Heat was the dominant driver of this physiological impairment, because in isolation it caused declines exceeding 45% in *P. astreoides* photophysiology, metabolism, symbiont density and growth compared to unheated treatments (Figure 3). In contrast, hypoxia alone produced limited physiological impairment yet magnified the negative impacts of heat across nearly all functional traits, resulting in responses that exceeded additive expectations.

Mechanistically, these synergistic effects likely arise from convergent disruption of mitochondrial and redox homeostasis: increased temperature destabilizes proteins, elevates metabolic demand, and enhances reactive oxygen species production (Helgoe et al., 2024b), while hypoxia limits terminal electron acceptors and disrupts redox balance (Dilworth et al., 2024; Gibbin et al., 2017; Seibel, 2011). At the molecular level, both stressors activate shared gene networks involving hypoxia-inducible factors (HIFα, EGLN1/PHD2) and oxidative stress responses (Alderdice, Hume, et al., 2022; Deleja et al., 2022; Zhang et al., 2023), indicating that oxygen limitation can intensify thermal- and light- induced stress and thus serve as a potentially fundamental, yet overlooked, contributor of coral bleaching.

Empirical evidence supports these mechanistic linkages between hypoxia and heat stress. For example, hypoxia (2 mg L^-1^) has been shown to lower thermal thresholds in *Acropora* sp. by 0.4–1.0 °C at temperatures exceeding 36 °C (Alderdice, Perna, et al., 2022) and increase mortality in *A. cervicornis* exposed to concurrent warming (Lucey et al., 2024), despite very short exposure durations (<24 hrs). Although species responses vary as taxa such as *Porites lutea* or *Monitpora tuberculosa* exhibited partial mitigation of photosynthetic declines under 9 days of combined heat and hypoxia (Jain et al. 2023), the overall pattern observed here suggests that shared energetic and redox constraints drive synergistic vulnerability. While these previous studies show relatively uniform responses consistent with our experiments, more standardized experiments are needed to broaden species representation and better resolve responses to combined stressors.

### Additive effects are prevalent under combined hypoxia and acidification

When hypoxia was compounded with acidification, additive effects were the most commonly occurring (n = 12), consistent with meta-analyses on non-coral taxa (Boyd et al., 2018; Steckbauer et al., 2020). The high number of additive outcomes observed under acidification, compared to heat with hypoxia, suggests that independent physiological pathways can regulate coral responses to low dissolved oxygen and pH at the tested intensities and durations.

In *S. siderea*, additive effects were detected across six measured parameters, including calcification and respiration – responses typically negatively affected by individual acidification and hypoxia stress, respectively (e.g.,Hughes et al., 2022; Leung et al., 2022). The only exceptions were a synergistic increase in photosynthetic rates and an antagonistic effect on biomass. This species exhibited complete resistance to acidification in the absence of other stressors, indicating that the additive outcomes largely reflected the physiological influence of hypoxia alone or its interaction with acidification. For *P. furcata*, additive effects were equally common but somewhat differed by trait (i.e., photosynthesis, P:R ratio, symbiont density, chlorophyll content, and photochemical efficiency, as well as biomass). The only exceptions in this species were two synergistic declines that exceeded additive expectations by 0.4% for respiration and 15.8% in calcification.

Although synergistic effects were also observed under combined hypoxia and acidification, these were fewer (n = 3) than those observed under combined hypoxia and heat (n = 7), with the overall prevalence of additive outcomes demonstrating mostly independent physiological pathways. However, such independence has limits, for example oxygen availability is necessary for energetically costly processes like calcification, which is also usually pH sensitive and typically affected over periods longer than a couple of weeks (e.g. Colombo-Pallotta et al., 2010; Holcomb et al., 2014, Leung et al. 2022). Thus, even additive stressor effects have the potential amplify long-term energetic deficits. To date, only two short-term studies (≤ 5 h) have tested the combined effects of hypoxia and acidification in corals, showing that dark calcification was strongly suppressed under hypoxia regardless of pH (Wijgerde et al., 2014), whereas neither net photosynthesis or dark respiration were affected by low DO or pH (*Galaxea fascicularis*; Osinga et al., 2017; though note this was also dependent on flow conditions). Given that hypoxia typically co-occurs with acidification due to stoichiometric coupling of O_2_ and CO_2_ (Gobler & Baumann, 2016), this warrant further investigation into the combined effects of oxygen limitation and acidification.

### Respiration as a sensitive indicator of stress

Among all measured traits, dark respiration was the most consistently suppressed across the three stressors, except for *S. siderea* under acidification alone, reinforcing its role as a sensitive and integrative indicator of coral physiological stress. Respiration reflects the reliance on oxygen for mitochondrial ATP production and maintenance of redox homeostasis (Handy & Loscalzo, 2012), providing a direct link into the coral energetic state under environmental perturbations. Metabolic depression under hypoxia is commonly observed and interpreted generally as an adaptive response that reduces ATP demand to conserve energy under oxygen limitation (Linsmayer et al., 2024; Murphy & Richmond, 2016). However, this strategy carries significant trade-offs, such as reductions in aerobic metabolism or shifts to anaerobic pathways that lower total ATP yield, constraining growth, calcification or cellular repair under dark conditions.

Our results, together with prior work, indicate that such depression may initially buffer short-term hypoxia exposure but becomes detrimental under prolonged stress. This is consistent with our 12–17 day experiments, where no bleaching or mortality occurred under < 2 mg O₂ L⁻¹, yet respiration and therefore also P:R ratios were highly affected. Nevertheless, P:R ratios remained greater than 1 demonstrating that net autotrophy was maintained despite these constraints. The species tested in the present study are comparatively hypoxia tolerant, as observations of hypoxia-sensitive species in other studies, which often lasted only a few hours or days, reported rapid bleaching, tissue sloughing, or mortality (Alderdice, Perna, et al., 2022; Deleja et al., 2022; Lucey et al., 2024).

Thermal stress can also depress respiration in many coral species (Parry et al., 2025), although transient increases may occur as metabolic demand rises during initial exposure (Lesser, 2006). Acidification alone rarely produces strong directional respiratory responses (e.g. Leung et al., 2022), yet meta-analytic evidence across ectotherms indicates that combined ocean acidification and warming generally produce additive or occasionally antagonistic effects, with the dominant driver shaping the response (Alter et al., 2024; Lefevre, 2016). Importantly, even when directional changes are absent, deviations from baseline respiration are widespread, reflecting energetic regulation and compensatory costs under moderate and high stress (Alter et al., 2024). Consequently, respiration suppression under combined stressors such as hypoxia, heat and acidification likely reflects both additive and synergistic energetic constraints, making it a sensitive early-warning signal of coral physiological compromise before bleaching or mortality occurs.

### Ecological implications and resilience of coral reefs under compound events

Our study advances understanding hypoxia-related compound stress on coral physiology through extended-exposure experiments, highlighting the strong potential for hypoxia to synergistically interact with other climate change stressors. While we examined only three, relatively stress-tolerant species, we view this work as an important starting point intended to catalyze broader research on how hypoxia interacts with multiple environmental stressors. Our findings are broadly aligned with meta-analyses investigating hypoxia-related compound stress in marine organisms (Steckbauer et al., 2020), supporting the relevance of our results beyond the specific species examined. We emphasize that additional multi-stressor experiments encompassing a wider diversity of coral taxa are urgently needed to evaluate the generality and ecological significance of these responses. Our results demonstrate that coral performance under climate change will be shaped not only by the intensity of individual stressors but increasingly by their timing and co-occurrence, including during extreme compound events.

Hypoxia, previously regarded as largely confined to temperate regions and uncommon on tropical reefs, is becoming more prevalent due to eutrophication, warming, and ocean stratification (Altieri & Gedan, 2015; Hughes et al., 2020), and has recently been shown to structure coral community composition (Lucey et al., 2024). While hypoxia alone primarily affected respiration, in combination with heat or acidification it produced stronger impacts on symbiosis health and growth, with both additive and synergistic outcomes, emphasizing the high variability in outcomes that warrant further mechanistic study.

Ecologically, our results underscore that hypoxia should be considered a central factor in reef futures (e.g., Deutsch et al., 2024; Pezner et al., 2023). While hypoxia alone may not cause severe physiological declines in resistant species, it can exacerbate the effects of heat and, to a lesser extent, acidification, on key traits, accelerating reef degradation under climate change. Disturbance-tolerant taxa may persist under summative stress, but with trade-offs including simplified community structure, reduced biodiversity, and altered ecosystem functioning (Lucey et al., 2024). More sensitive taxa are likely to decline more rapidly, and even resilient “winner” species will face limits under increasingly severe marine heatwaves. Assemblages dominated by tolerant species can continue to support reef accretion, though in modified ecological and structural forms, consistent with historical records of non-acroporid Caribbean reefs maintaining slow but steady growth through past perturbations (Islas-Dominguez et al., 2025). Ultimately, reef persistence will depend on species-specific capacities to withstand the metabolic, ecological, and evolutionary consequences of escalating compound stress. Integrated climate mitigation and local management of nutrients are needed to ensure reefs remain within the safe operating space of planetary boundaries and avoid threshold-crossing events or metabolic storms that could push these systems toward collapse.

## Acknowledgements

We thank the staff of CARMABI and STRI for logistical support, Stacy Shinneman for map preparation, Ilmer Benda for laboratory analyses, and Bas van Beusekom for assistance with experimental setup. Funding for RvO was provided by the Volkert van der Willigen Fonds of the Stichting Amsterdamse Universiteits Fonds. The remainder of the project was supported by a VIDI grant of the Dutch Research Council (NWO) awarded to VS (VI.Vidi.203.069).

## Supplemental Information

### Supplementary Materials and Methods

#### Experiment 1: Hypoxia × acidification

Temperature was maintained using an aquarium controller (Apex Pro, Neptune Systems, USA) with probes accurate to ±0.1°C, heaters (ThermoControl 200–300, EHEIM, DEU), temperature loggers (ENVloggers T7.3, Electric Blue, sampling interval 10 min, precision ≤ 0.1 °C, accuracy ≤ 0.2 °C), and circulation pumps (Nano Voyager 1000 L h⁻¹, Sicce, ITA).

Oxygen concentrations were maintained in each independent tank by bubbling seawater with nitrogen gas or ambient air, with gas injection controlled through a DO feedback and solenoid valves (Neptune Systems). Oxygen levels were monitored every minute in each tank by an OxyGuard DO probe connected to an aquarium controller (Neptune Systems, Apex Aquacontroller), which opened or closed the respective solenoid valves to add nitrogen or air to maintain the programmed treatment conditions. Oxygen probes were calibrated at the start of experiments following the manufacturer’s protocol, and frequently checked against a handheld multimeter. Tanks were fitted with acrylic lids to stabilize oxygen concentrations.

All tanks were equipped with pH probes (Neptune Systems, USA) and gas lines supplying either CO₂ or CO₂-free air (scrubbed via a soda lime cylinder) to regulate pH and counteract artificial increases caused by N₂ bubbling. Acidic treatments were maintained on the total pH scale (pH_T_) by applying an offset to the Neptune pH probes, frequently adjusted against pH_T_ values measured with a handheld mV electrode (PHC101 HQ40d, HACH, USA) and standardized by measuring TRIS buffer purchased from A. Dickson (Scripps) at multiple temperatures (Paulsen & Dickson, 2020).

Each tank received seawater from a reservoir replenished twice daily with 0.5 μm-filtered seawater (Melt Blown Polypropylene Filter, H2O Filter Warehouse, USA) pumped from 10 m depth at a renewal rate of 26.3 L day⁻¹. Coral positions were rotated daily, and corals were fed five times during the experiment with live *Artemia* (50 mL, resulting tank concentration of ∼0.35 individuals mL⁻¹), with circulation pumps and gas inputs turned off for 30 min during feeding.

Discrete water samples (n = 5 per tank) were filtered (0.2 μm PES filters) into HDPE bottles, preserved with mercuric chloride (0.02% of sample volume), and stored at 4°C until analysis. Total alkalinity (TA) was measured via open-cell titration using a 716 DMS Titrino (Metrohm) following Dickson et al. (2007). Certified reference material verified accuracy (certified value: 2202.75 ± 0.70 μmol·kg⁻¹; measured values: 2202 ± 10 μmol·kg⁻¹, mean ± SD, n = 14).

Photosynthetically active radiation (PAR) followed a simulated diel cycle using LED lights (CoralCare Gen II, Philips, NLD): ramp-up from 05:30 to 07:00, peak intensity of 376 ± 28 μmol photons m⁻² s⁻¹ (*n* = 95) between 07:00 and 17:00, and ramp-down from 17:00 to 18:30.

For dark-adapted maximum quantum yield of photosystem II (F_v_/F_m_), PAM settings included a measuring light of 6 on days 1–3 and 8 on days 5–15, with saturation intensity set to 6, saturation width to 0.86 s, gain to 1–2, and damping to 2 (red-light pulse amplitude modulated fluorometer; Diving-PAM-I, Walz GmbH, Germany).

#### Experiment 2: Hypoxia × heat

Methods are described in detail only for procedures that differed from those used in Experiment 1. Water circulation within the tanks was achieved with submersible circulation pumps (Aquaneat Wavemaker, 800 GPH). Daily measurements of temperature, pHT, and salinity were conducted in all tanks using a multiparameter meter (HQ40d, HACH, USA), with DO monitored through the Apex OxyGuard DO probes. Each tank received seawater pumped from 3 m depth and filtered using a Bubble Bead XF 20,000 system (Propulsion Pools, MYS), with a tank replenishment rate of 96 L day⁻¹. Discrete water samples (*n* = 3 per tank) were collected throughout the experiment to measure TA using the same method as in Experiment 1.

Tanks were illuminated with photosynthetically active radiation (PAR) following a 12:12 h light:dark cycle using LED lights (Hydra 64 HD, Aqua Illumination, USA): ramp-up from 05:00 to 06:00, peak intensity of 280 ± 29 μmol photons m⁻² s⁻¹ (*n* = 114) between 06:00 and 16:00, and ramp-down from 16:00 to 17:00. Light was measured using a PAR sensor (MINI-SPEC for Diving-PAM II, Walz GmbH, DEU), and coral positions rotated daily. For measuring F_v_/F_m_, the measuring light intensity was set to 11, saturation intensity to 4, saturation width to 8, gain to 1–2, damping to 2, and frequency to 3 (red-light pulse amplitude modulated fluorometer; Diving-PAM-II, Walz GmbH, Germany).

### Supplementary Tables

**SI Table 1.** Treatment conditions for Experiments 1 and 2 across tank replicates, with sample sizes (*n*) provided. Temperature was measured continuously at 10-minute intervals.

| Experiment | Treatment | Value | T<br>(°C) | pH<sub>T</sub><br>(total<br>scale) | Dissolved<br>oxygen<br>(mg L<sup>-1</sup>) | Salinity<br>(ppt) | TA<br>(μmol<br>kg<sup>-1</sup>) |
| --- | --- | --- | --- | --- | --- | --- | --- |
| 1 | Control | mean | 27.6 | 7.94 | 6.25 | 35.2 | 2195 |
|  |  | SD | 0.29 | 0.07 | 0.33 | 0.14 | 51 |
|  |  | n | 5358 | 36 | 36 | 34 | 10 |
|  | Hypoxic | mean | 27.6 | 7.99 | 1.95 | 35.2 | 2189 |
|  |  | SD | 0.28 | 0.07 | 0.44 | 0.23 | 73 |
|  |  | n | 5196 | 36 | 36 | 32 | 10 |
|  | Acidic | mean | 27.5 | 7.67 | 6.84 | 35.2 | 2224 |
|  |  | SD | 0.34 | 0.07 | 0.47 | 0.23 | 99 |
|  |  | n | 5480 | 36 | 36 | 34 | 10 |
|  | Hyp:Acid | mean | 27.5 | 7.66 | 1.96 | 35.1 | 2242 |
|  |  | SD | 0.29 | 0.07 | 0.45 | 0.15 | 69 |
|  |  | n | 5196 | 36 | 36 | 32 | 10 |
| 2 | Control | mean | 28.1 | 7.94 | 6.20 | 32.8 | 2066 |
|  |  | SD | 0.18 | 0.08 | 0.53 | 0.35 | 87 |
|  |  | n | 30 | 41 | 1303 | 44 | 6 |
|  | Hypoxic | mean | 28.2 | 7.96 | 1.27 | 32.9 | 2113 |
|  |  | SD | 0.19 | 0.06 | 0.20 | 0.42 | 55 |
|  |  | n | 30 | 40 | 32398 | 44 | 6 |
|  | Hot | mean | 32.3 | 7.93 | 5.40 | 33.3 | 2209 |
|  |  | SD | .3 | 0.09 | 0.44 | 0.46 | 19 |
|  |  | n | 30 | 39 | 33672 | 44 | 6 |
|  | Hyp:Hot | mean | 32.2 | 7.95 | 1.32 | 33.2 | 2183 |
|  |  | SD | 0.33 | 0.06 | 0.21 | 0.47 | 11 |
|  |  | n | 30 | 40 | 31383 | 44 | 6 |

**SI Table 2.**
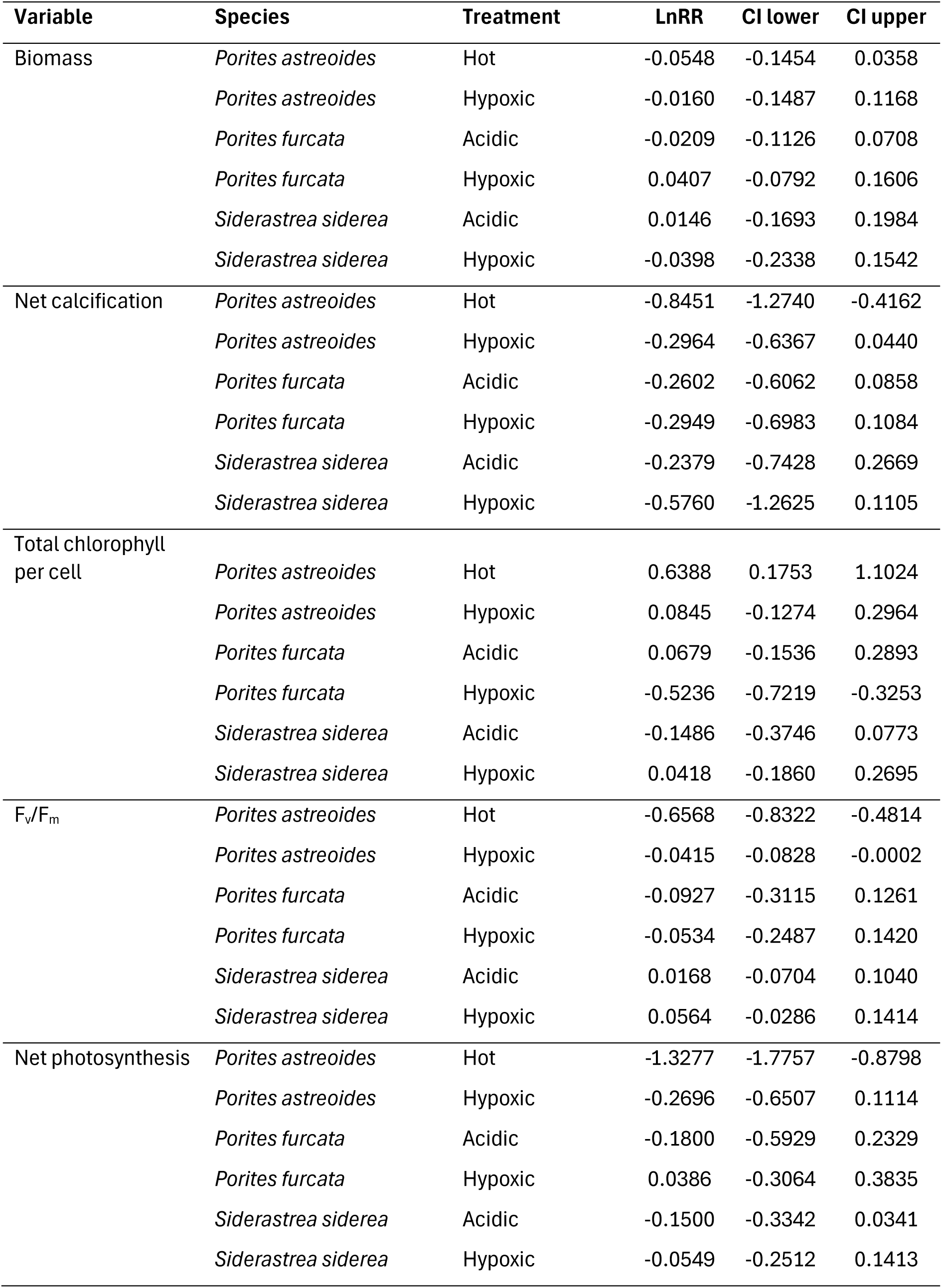
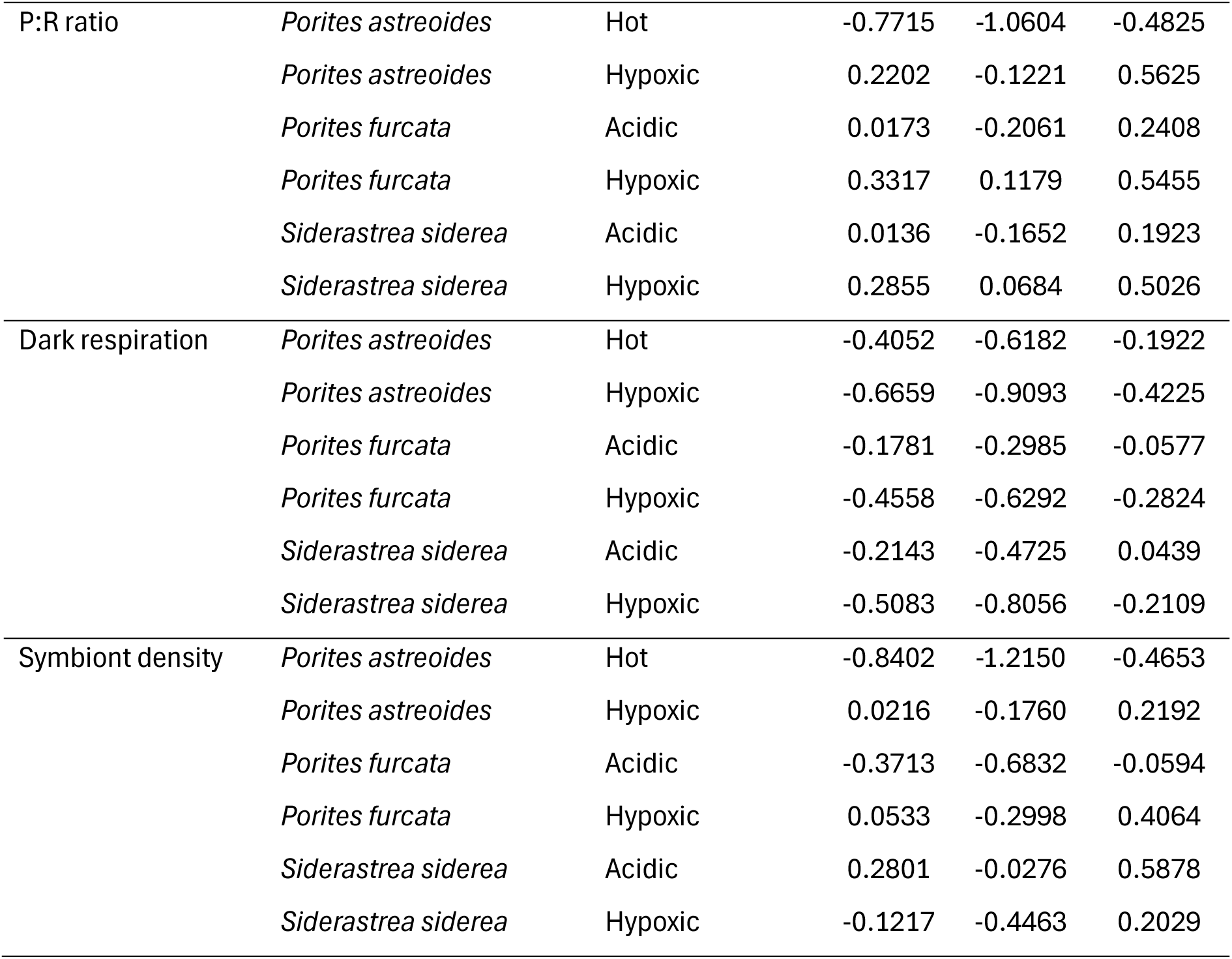
Independent stressor natural log response ratios (LnRR) and confidence interval (CI) values by physiological trait, species, and treatment.

**SI Table 3.**
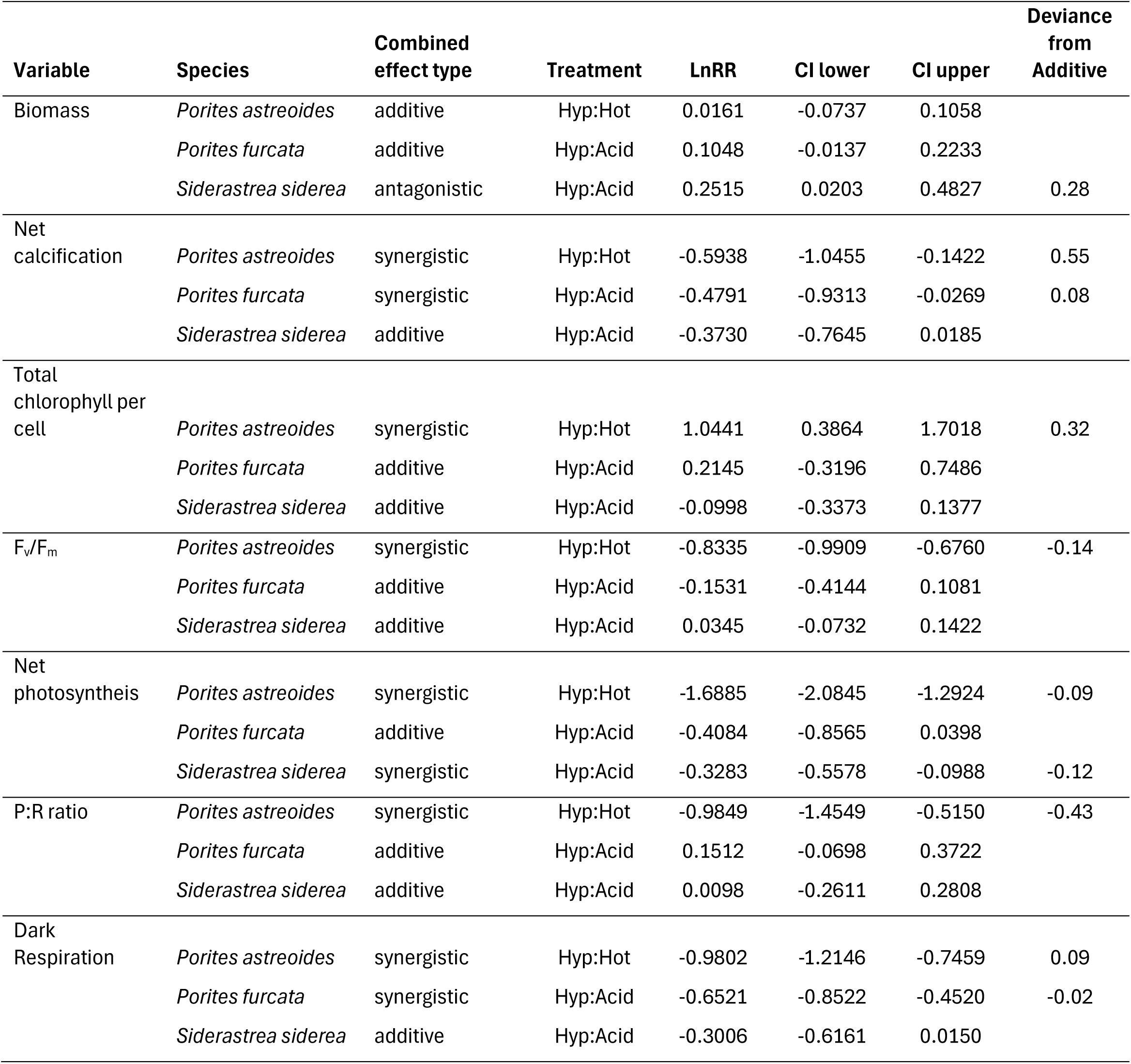
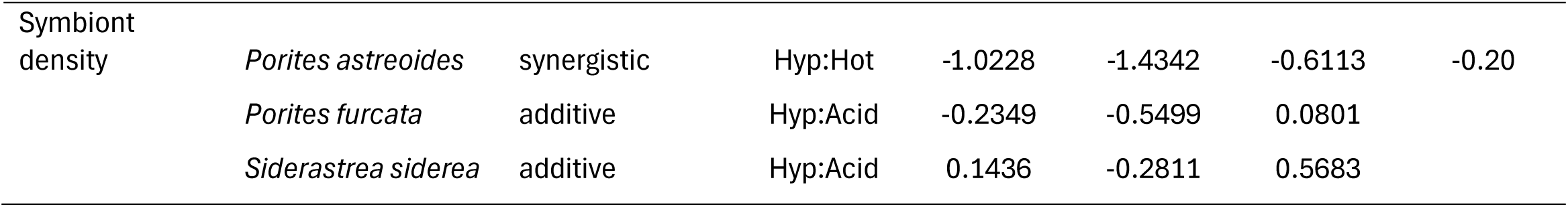
Combined stressor natural log response ratios (LnRR) and confidence interval (CI) values by physiological trait, species, categorized interaction type (based on the combined LnRR and the signs of the individual stressor LnRRs), and treatment. The last column shows the deviation from the calculated additive response, *Deviation_Additive_calc__*, which describes the magnitude of response beyond what is expected under an additive model.

### Supplementary Figures

**SI Figure 1.**
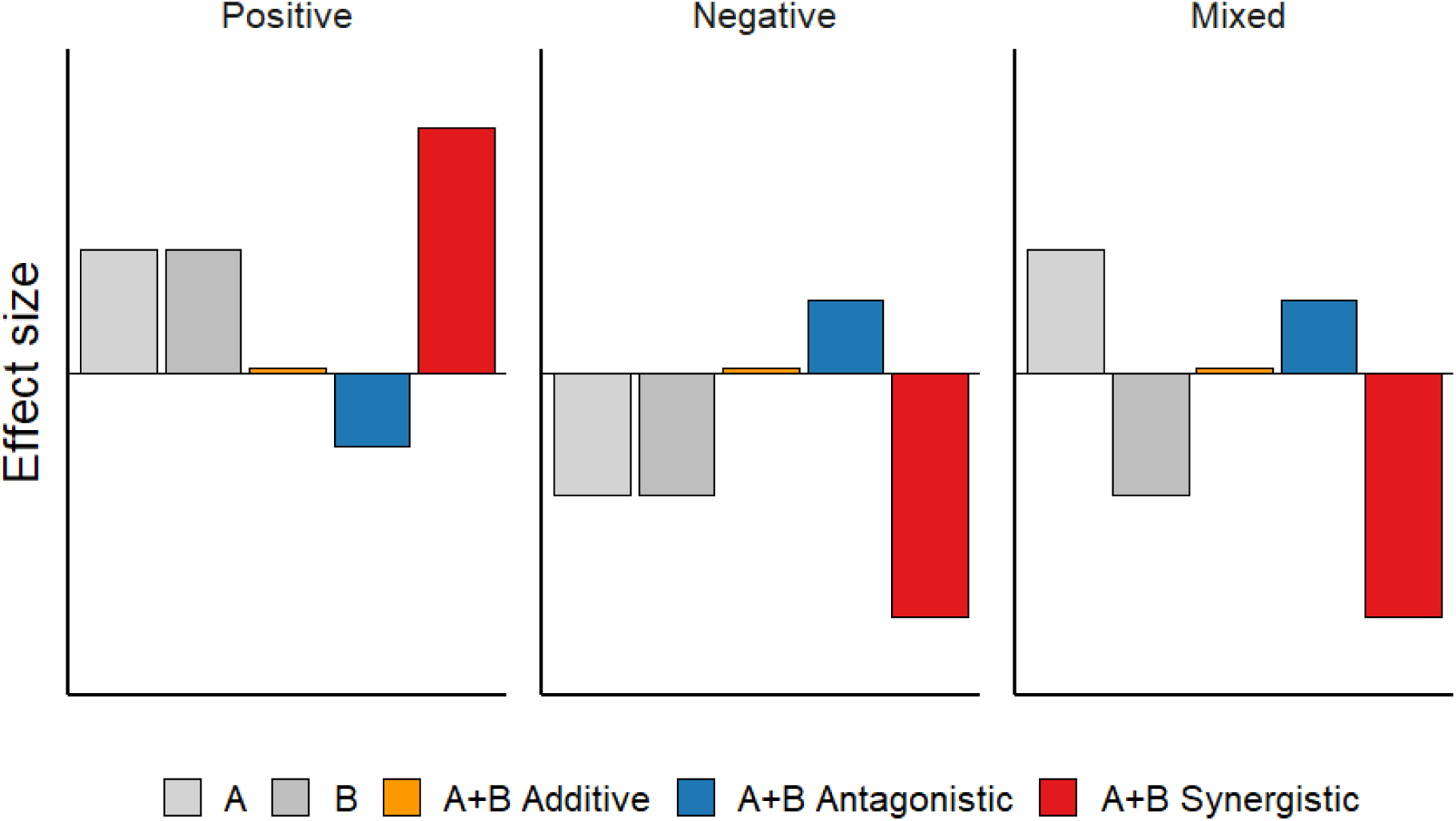
Conceptual illustration of combined stressor effects shown for three contexts: positive, negative, and mixed individual stressor effects. Bars represent individual stressors (gray tones), the additive outcome (orange), and observed interaction outcomes classified as antagonistic (blue) or synergistic (red). The additive bar overlaps with zero, indicating no deviation from the summation of the two effects. This schematic omits variance and confidence intervals for clarity and is intended for illustrative purposes only; in formal analyses, means from the control group are incorporated into effect size calculations, but these are not shown here. The positive combination is modeled after Gunderson et al. (2016)

**SI Figure 2.**
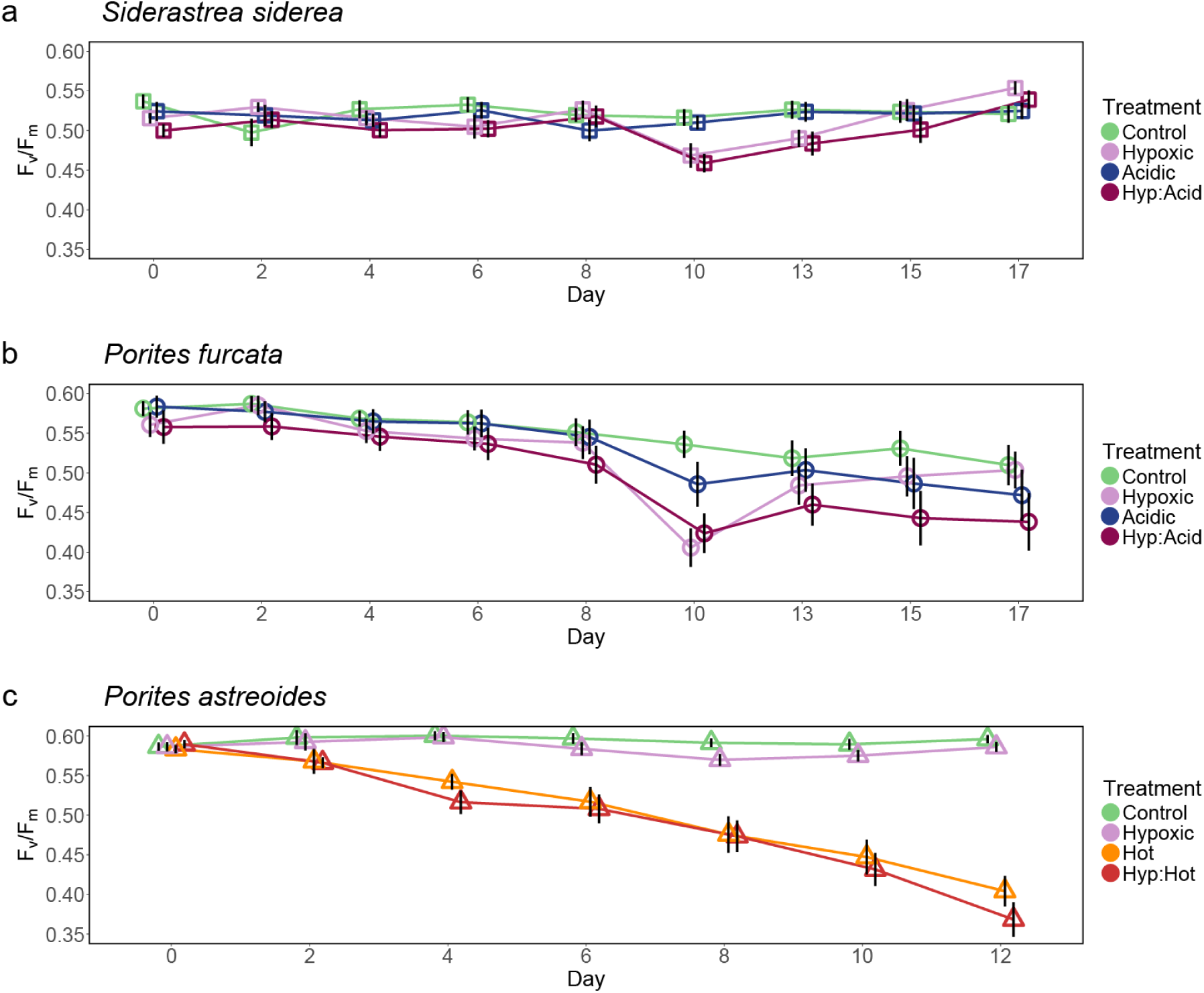
Maximum quantum yield of photosystem II (Fv/Fm) over time for three coral species in Experiments 1 and 2. Data represent treatment means ± SE of a) *Siderastrea siderea*, b) *Porites furcata*, and c) *Porites astreoides*. Note that the experiment for *P. astreoides* in Experiment 2 was five days shorter than Experiment 1.

## Literature Cited

Alderdice, R., Hume, B. C. C., Kühl, M., Pernice, M., Suggett, D. J., & Voolstra, C. R. (2022). Disparate Inventories of Hypoxia Gene Sets Across Corals Align With Inferred Environmental Resilience. Frontiers in Marine Science, 9. 10.3389/FMARS.2022.834332

Alderdice, R., Perna, G., Cárdenas, A., Hume, B. C. C., Wolf, M., Kühl, M., Pernice, M., Suggett, D. J., & Voolstra, C. R. (2022). Deoxygenation lowers the thermal threshold of coral bleaching. Scientific Reports, 12(1). 10.1038/s41598-022-22604-3

Alter, K., Jacquemont, J., Claudet, J., Lattuca, M. E., Barrantes, M. E., Marras, S., Manríquez, P. H., González, C. P., Fernández, D. A., Peck, M. A., Cattano, C., Milazzo, M., Mark, F. C., & Domenici, P. (2024). Hidden impacts of ocean warming and acidification on biological responses of marine animals revealed through meta-analysis. Nature Communications, 15(1), 1–13. 10.1038/S41467-024-47064-3;TECHMETA

Altieri, A. H., & Gedan, K. B. (2015). Climate change and dead zones. Global Change Biology, 21(4), 1395–1406. 10.1111/GCB.12754

Altieri, A. H., Harrison, S. B., Seemann, J., Collin, R., Diaz, R. J., & Knowlton, N. (2017a). Tropical dead zones and mass mortalities on coral reefs. Proceedings of the National Academy of Sciences of the United States of America, 114(14), 3660–3665. 10.1073/pnas.1621517114

Altieri, A. H., Harrison, S. B., Seemann, J., Collin, R., Diaz, R. J., & Knowlton, N. (2017b). Tropical dead zones and mass mortalities on coral reefs. Proceedings of the National Academy of Sciences of the United States of America, 114(14), 3660–3665. 10.1073/PNAS.1621517114/SUPPL_FILE/PNAS.201621517SI.PDF

Armstrong McKay, D. I., Staal, A., Abrams, J. F., Winkelmann, R., Sakschewski, B., Loriani, S., Fetzer, I., Cornell, S. E., Rockström, J., & Lenton, T. M. (2022). Exceeding 1.5°C global warming could trigger multiple climate tipping points. Science, 377(6611). 10.1126/SCIENCE.ABN7950/SUPPL_FILE/SCIENCE.ABN7950_DATA_S1.ZIP

Boyd, P. W., Collins, S., Dupont, S., Fabricius, K., Gattuso, J. P., Havenhand, J., Hutchins, D. A., Riebesell, U., Rintoul, M. S., Vichi, M., Biswas, H., Ciotti, A., Gao, K., Gehlen, M., Hurd, C. L., Kurihara, H., McGraw, C. M., Navarro, J. M., Nilsson, G. E., … Pörtner, H. O. (2018). Experimental strategies to assess the biological ramifications of multiple drivers of global ocean change—A review. Global Change Biology, 24(6), 2239–2261. 10.1111/GCB.14102

Breitburg, D., Levin, L. A., Oschlies, A., Grégoire, M., Chavez, F. P., Conley, D. J., Garçon, V., Gilbert, D., Gutiérrez, D., Isensee, K., Jacinto, G. S., Limburg, K. E., Montes, I., Naqvi, S. W. A., Pitcher, G. C., Rabalais, N. N., Roman, M. R., Rose, K. A., Seibel, B. A., … Zhang, J. (2018). Declining oxygen in the global ocean and coastal waters. Science, 359(6371). 10.1126/SCIENCE.AAM7240

Burger, F. A., Terhaar, J., & Frölicher, T. L. (2022). Compound marine heatwaves and ocean acidity extremes. Nature Communications 2022 13:1, 13(1), 1–12. 10.1038/s41467-022-32120-7

Colombo-Pallotta, M. F., Rodríguez-Román, A., & Iglesias-Prieto, R. (2010). Calcification in bleached and unbleached Montastraea faveolata: Evaluating the role of oxygen and glycerol. Coral Reefs, 29(4), 899–907. 10.1007/S00338-010-0638-X/METRICS

Crain, C. M., Kroeker, K., & Halpern, B. S. (2008). Interactive and cumulative effects of multiple human stressors in marine systems. Ecology Letters, 11(12), 1304–1315. 10.1111/J.1461-0248.2008.01253.X

Deleja, M., Paula, J. R., Repolho, T., Franzitta, M., Baptista, M., Lopes, V., Simão, S., Fonseca, V. F., Duarte, B., & Rosa, R. (2022). Effects of Hypoxia on Coral Photobiology and Oxidative Stress. Biology, 11(7). 10.3390/biology11071068

Deutsch, C., Penn, J. L., & Lucey, N. (2024). Climate, Oxygen, and the Future of Marine Biodiversity. Annual Review of Marine Science, 16, 217–245. 10.1146/annurev-marine-040323-095231

Diaz, R. J., & Rosenberg, R. (2008). Spreading dead zones and consequences for marine ecosystems. Science, 321(5891), 926–929. 10.1126/SCIENCE.1156401/SUPPL_FILE/1156401-DIAZ-SOM.PDF

Dilworth, J., Million, W. C., Ruggeri, M., Hall, E. R., Dungan, A. M., Muller, E. M., & Kenkel, C. D. (2024). Synergistic response to climate stressors in coral is associated with genotypic variation in baseline expression. Proceedings of the Royal Society B: Biological Sciences, 291(2019), 20232447. 10.1098/RSPB.2023.2447

Ferrer, E. M., Pezner, A. K., Eddebbar, Y. A., Breitburg, D., Crowe, S., Garçon, V., Grégoire, M., Jane, S. F., Leavitt, P. R., Levin, L., Rose, K., & Wallace, D. (2025). Why aquatic deoxygenation belongs in the planetary boundary framework. PLOS Climate, 4(5), e0000619. 10.1371/JOURNAL.PCLM.0000619

Fitt, W. K., McFarland, F. K., Warner, M. E., & Chilcoat, G. C. (2000). Seasonal patterns of tissue biomass and densities of symbiotic dinoflagellates in reef corals and relation to coral bleaching. Limnology and Oceanography, 45(3), 677–685. 10.4319/LO.2000.45.3.0677;PAGE:STRING:ARTICLE/CHAPTER

Gedan, K. B., Altieri, A. H., Feller, I., Burrel, R., & Breitburg, D. (2017). Community composition in mangrove ponds with pulsed hypoxic and acidified conditions. Ecosphere, 8(12). 10.1002/ecs2.2053

Gibbin, E. M., Chakravarti, L. J., Jarrold, M. D., Christen, F., Turpin, V., N’Siala, G. M., Blier, P. U., & Calosi, P. (2017). Can multi-generational exposure to ocean warming and acidification lead to the adaptation of life history and physiology in a marine metazoan? Journal of Experimental Biology, 220(4), 551–563. 10.1242/JEB.149989/262461/AM/CAN-MULTI-GENERATIONAL-EXPOSURE-TO-OCEAN-WARMING

Gobler, C. J., & Baumann, H. (2016). Hypoxia and acidification in ocean ecosystems: coupled dynamics and effects on marine life. Biology Letters, 12(5). 10.1098/RSBL.2015.0976

Gruber, N., Boyd, P. W., Frölicher, T. L., & Vogt, M. (2021). Biogeochemical extremes and compound events in the ocean. Nature 2021 600:7889, 600(7889), 395–407. 10.1038/s41586-021-03981-7

Gunderson, A. R., Armstrong, E. J., & Stillman, J. H. (2016). Multiple Stressors in a Changing World: The Need for an Improved Perspective on Physiological Responses to the Dynamic Marine Environment. https://doi.org/10.1146/Annurev-Marine-122414-033953, 8, 357–378. 10.1146/ANNUREV-MARINE-122414-033953

Gurevitch, J., Morrison, J. A., & Hedges, L. V. (2000). The interaction between competition and predation: A meta-analysis of field experiments. American Naturalist, 155(4), 435–453. 10.1086/303337,

Haas, A. F., Smith, J. E., Thompson, M., & Deheyn, D. D. (2014). Effects of reduced dissolved oxygen concentrations on physiology and fluorescence of hermatypic corals and benthic algae. PeerJ, 2014(1). 10.7717/peerj.235

Handy, D. E., & Loscalzo, J. (2012). Redox Regulation of Mitochondrial Function. Antioxidants & Redox Signaling, 16(11), 1323. 10.1089/ARS.2011.4123

Harvey, B. P., Gwynn-Jones, D., & Moore, P. J. (2013). Meta-analysis reveals complex marine biological responses to the interactive effects of ocean acidification and warming. Ecology and Evolution, 3(4), 1016–1030. 10.1002/ECE3.516

Hedges, L. V., Gurevitch, J., & Curtis, P. S. (1999). THE META-ANALYSIS OF RESPONSE RATIOS IN EXPERIMENTAL ECOLOGY. Ecology, 80(4), 1150–1156. 10.1890/0012-9658(1999)080[1150:TMAORR]2.0.CO;2

Helgoe, J., Davy, S. K., Weis, V. M., & Rodriguez-Lanetty, M. (2024a). Triggers, cascades, and endpoints: connecting the dots of coral bleaching mechanisms. Biological Reviews, 000–000. 10.1111/BRV.13042

Helgoe, J., Davy, S. K., Weis, V. M., & Rodriguez-Lanetty, M. (2024b). Triggers, cascades, and endpoints: connecting the dots of coral bleaching mechanisms. Biological Reviews, 99(3), 715–752. 10.1111/BRV.13042

Holcomb, M., Tambutté, E., Allemand, D., & Tambutté, S. (2014). Light enhanced calcification in Stylophora pistillata: Effects of glucose, glycerol and oxygen. PeerJ, 2014(1), e375. 10.7717/PEERJ.375/SUPP-1

Hughes, D. J., Alderdice, R., Cooney, C., Kühl, M., Pernice, M., Voolstra, C. R., & Suggett, D. J. (2020). Coral reef survival under accelerating ocean deoxygenation. Nature Climate Change, 10(4), 296–307. 10.1038/s41558-020-0737-9

Hughes, D. J., Alexander, J., Cobbs, G., Kühl, M., Cooney, C., Pernice, M., Varkey, D., Voolstra, C. R., & Suggett, D. J. (2022). Widespread oxyregulation in tropical corals under hypoxia. Marine Pollution Bulletin, 179(May), 113722. 10.1016/j.marpolbul.2022.113722

Islas-Dominguez, E., Gischler, E., & Hudson, J. H. (2025). Holocene development of submerged keep-up patch reefs on Bermuda without acroporids: A model of future reef accretion. The Depositional Record. 10.1002/DEP2.70023

Jackson, M. C., Loewen, C. J. G., Vinebrooke, R. D., & Chimimba, C. T. (2016). Net effects of multiple stressors in freshwater ecosystems: a meta-analysis. Global Change Biology, 22(1), 180–189. 10.1111/GCB.13028

Jain, T., Buapet, P., Ying, L., & Yucharoen, M. (2023). Differing Responses of Three Scleractinian Corals from Phuket Coast in the Andaman Sea to Experimental Warming and Hypoxia. Journal of Marine Science and Engineering, 11(2). 10.3390/jmse11020403

Jeffrey, S. W., & Humphrey, G. F. (1975). New spectrophotometric equations for determining chlorophylls a, b, c1 and c2 in higher plants, algae and natural phytoplankton. Biochemie Und Physiologie Der Pflanzen, 167(2), 191–194. 10.1016/S0015-3796(17)30778-3

Johnson, M. D., Scott, J. J., Leray, M., Lucey, N., Bravo, L. M. R., Wied, W. L., & Altieri, A. H. (2021). Rapid ecosystem-scale consequences of acute deoxygenation on a Caribbean coral reef. Nature Communications, 12(1), 1–12. 10.1038/s41467-021-24777-3

Johnson, M. D., Swaminathan, S. D., Nixon, E. N., Paul, V. J., & Altieri, A. H. (2021). Differential susceptibility of reef-building corals to deoxygenation reveals remarkable hypoxia tolerance. Scientific Reports, 11(1). 10.1038/s41598-021-01078-9

Jokiel, P. L., Maragos, J. E., & Franzisket, L. (1978). Coral growth: buoyant weight technique. In D. R. Stoddart & R. E. Johannes (Eds.), Coral Reefs: research methods (Vol. 5, pp. 529–541). Unesco. https://unesdoc.unesco.org/ark:/48223/pf0000029306.locale=en

Keeling, R. F., Körtzinger, A., & Gruber, N. (2009). Ocean Deoxygenation in a Warming World. https://doi.org/10.1146/Annurev.Marine.010908.163855, 2(1), 199–229. 10.1146/ANNUREV.MARINE.010908.163855

Kvitt, H., Malik, A., Ben-Tabou de-Leon, S., Shemesh, E., Lalzar, M., Gruber, D. F., Rosenfeld, H., Shi, T., Mass, T., & Tchernov, D. (2022). Transcriptional responses indicate acclimation to prolonged deoxygenation in the coral Stylophora pistillata. Frontiers in Marine Science, 9. 10.3389/fmars.2022.999558

Lavy, A., Eyal, G., Neal, B., Keren, R., Loya, Y., & Ilan, M. (2015). A quick, easy and non-intrusive method for underwater volume and surface area evaluation of benthic organisms by 3D computer modelling. Methods in Ecology and Evolution, 6(5), 521–531. 10.1111/2041-210X.12331;PAGEGROUP:STRING:PUBLICATION

Lefevre, S. (2016). Are global warming and ocean acidification conspiring against marine ectotherms? A meta-analysis of the respiratory effects of elevated temperature, high CO2 and their interaction. Conservation Physiology, 4(1). 10.1093/CONPHYS/COW009

Lesser, M. P. (2006). Oxidative stress in marine environments: Biochemistry and physiological ecology. Annual Review of Physiology, 68(Volume 68, 2006), 253–278. 10.1146/ANNUREV.PHYSIOL.68.040104.110001/CITE/REFWORKS

Leung, J. Y. S., Zhang, S., Connell, S. D., Leung, J. Y. S., Zhang, S., & Connell, S. D. (2022). Is Ocean Acidification Really a Threat to Marine Calcifiers? A Systematic Review and Meta-Analysis of 980+ Studies Spanning Two Decades. Small, 18(35), 2107407. 10.1002/SMLL.202107407

Linsmayer, L. B., Deheyn, D. D., Tomanek, L., & Tresguerres, M. (2020). Dynamic regulation of coral energy metabolism throughout the diel cycle. Scientific Reports 2020 10:1, 10(1), 1–11. 10.1038/s41598-020-76828-2

Linsmayer, L. B., Noel, S. K., Leray, M., Wangpraseurt, D., Hassibi, C., Kline, D. I., & Tresguerres, M. (2024). Effects of bleaching on oxygen dynamics and energy metabolism of two Caribbean coral species. Science of The Total Environment, 919, 170753. 10.1016/J.SCITOTENV.2024.170753

Lucey, N. M., César-Ávila, C., Eckert, A., Rajagopalan, A., Brister, W. C., Kline, E., Altieri, A. H., Deutsch, C. A., & Collin, R. (2024). Coral Community Composition Linked to Hypoxia Exposure. Global Change Biology, 30(10). 10.1111/gcb.17545

Mallon, J. E., Altieri, A. H., Cyronak, T., Melendez-Declet, C. V., Paul, V. J., & Johnson, M. D. (2025). Sublethal changes to coral metabolism in response to deoxygenation. Journal of Experimental Biology, 228(4). 10.1242/JEB.249638/365403/AM/SUBLETHAL-CHANGES-TO-CORAL-METABOLISM-IN-RESPONSE

Murphy, J. W. A., & Richmond, R. H. (2016). Changes to coral health and metabolic activity under oxygen deprivation. PeerJ, 2016(4). 10.7717/peerj.1956

Nakagawa, S., & Cuthill, I. C. (2007). Effect size, confidence interval and statistical significance: a practical guide for biologists. Biological Reviews, 82(4), 591–605. 10.1111/J.1469-185X.2007.00027.X

Nakagawa, S., & Schielzeth, H. (2013). A general and simple method for obtaining R2 from generalized linear mixed-effects models. Methods in Ecology and Evolution, 4(2), 133–142. 10.1111/J.2041-210X.2012.00261.X;SUBPAGE:STRING:FULL

Nelson, H. R., & Altieri, A. H. (2019). Oxygen: the universal currency on coral reefs. Coral Reefs 2019 38:2, 38(2), 177–198. 10.1007/S00338-019-01765-0

Osinga, R., Derksen-Hooijberg, M., Wijgerde, T., & Verreth, J. A. J. (2017). Interactive effects of oxygen, carbon dioxide and flow on photosynthesis and respiration in the scleractinian coral Galaxea fascicularis. Journal of Experimental Biology, 220(12), 2236–2242. 10.1242/JEB.140509/262346/AM/INTERACTIVE-EFFECTS-OF-OXYGEN-CARBON-DIOXIDE-AND

Parry, A. J., Klein, S. G., & Duarte, C. M. (2025). Thermal extremes likely trigger metabolic imbalance in coral holobionts. Scientific Reports, 15(1), 1–12. 10.1038/S41598-025-16880-Y;SUBJMETA

Pezner, A. K., Courtney, T. A., Barkley, H. C., Chou, W.-C., Chu, H.-C., Clements, S. M., Cyronak, T., DeGrandpre, M. D., Kekuewa, S. A. H., Kline, D. I., Liang, Y.-B., Martz, T. R., Mitarai, S., Page, H. N., Rintoul, M. S., Smith, J. E., Soong, K., Takeshita, Y., Tresguerres, M., … Andersson, A. J. (2023). Increasing hypoxia on global coral reefs under ocean warming. Nature Climate Change 2023, 1–7. 10.1038/s41558-023-01619-2

Rockström, J., Steffen, W., Noone, K., Persson, Å., Chapin, F. S., Lambin, E. F., Lenton, T. M., Scheffer, M., Folke, C., Schellnhuber, H. J., Nykvist, B., De Wit, C. A., Hughes, T., Van Der Leeuw, S., Rodhe, H., Sörlin, S., Snyder, P. K., Costanza, R., Svedin, U., … Foley, J. A. (2009). A safe operating space for humanity. Nature, 461(7263), 472–475. 10.1038/461472A;KWRD=SCIENCE

Sampaio, E., Santos, C., Rosa, I. C., Ferreira, V., Pörtner, H. O., Duarte, C. M., Levin, L. A., & Rosa, R. (2021). Impacts of hypoxic events surpass those of future ocean warming and acidification. Nature Ecology and Evolution, 5(3), 311–321. 10.1038/s41559-020-01370-3

Schneider, C. A., Rasband, W. S., & Eliceiri, K. W. (2012). NIH Image to ImageJ: 25 years of image analysis. Nature Methods, 9(7), 671–675. 10.1038/NMETH.2089,

Seibel, B. A. (2011). Critical oxygen levels and metabolic suppression in oceanic oxygen minimum zones. Journal of Experimental Biology, 214(2), 326–336. 10.1242/JEB.049171

Solomon, S. L., Lippens, C. R., Mazza, R., Powell, M. E., Johnson, K. W., Suvacarov, S., Zande, R. M. van der, & Schoepf, V. (2025). Physiological mechanisms underlying coral acclimatization capacity to novel, multi-stressor conditions. BioRxiv, 2025.05.08.652802. 10.1101/2025.05.08.652802

Steckbauer, A., Klein, S. G., & Duarte, C. M. (2020). Additive impacts of deoxygenation and acidification threaten marine biota. Global Change Biology, 26(10), 5602–5612. 10.1111/gcb.15252

Stramma, L., Schmidtko, S., Levin, L. A., & Johnson, G. C. (2010). Ocean oxygen minima expansions and their biological impacts. Deep Sea Research Part I: Oceanographic Research Papers, 57(4), 587–595. 10.1016/J.DSR.2010.01.005

Todgham, A. E., & Stillman, J. H. (2013). Physiological Responses to Shifts in Multiple Environmental Stressors: Relevance in a Changing World. Integrative and Comparative Biology, 53(4), 539–544. 10.1093/ICB/ICT086

Vaquer-Sunyer, R., & Duarte, C. M. (2011). Temperature effects on oxygen thresholds for hypoxia in marine benthic organisms. Global Change Biology, 17(5), 1788–1797. 10.1111/j.1365-2486.2010.02343.x

Wijgerde, T., Silva, C. I. F., Scherders, V., Van Bleijswijk, J., & Osinga, R. (2014). Coral calcification under daily oxygen saturation and pH dynamics reveals the important role of oxygen. Biology Open, 3(6), 489–493. 10.1242/bio.20147922

Woods, H. A., Moran, A. L., Atkinson, D., Audzijonyte, A., Berenbrink, M., Borges, F. O., Burnett, K. G., Burnett, L. E., Coates, C. J., Collin, R., Costa-Paiva, E. M., Duncan, M. I., Ern, R., Laetz, E. M. J., Levin, L. A., Lindmark, M., Lucey, N. M., McCormick, L. R., Pierson, J. J., … Verberk, W. C. E. P. (2022). Integrative Approaches to Understanding Organismal Responses to Aquatic Deoxygenation. Biological Bulletin, 243(2), 85–103. 10.1086/722899/ASSET/IMAGES/LARGE/FG3.JPEG

Zhang, K., Wu, Z., Liu, Z., Tang, J., Cai, W., An, M., & Zhou, Z. (2023). Acute hypoxia induces reduction of algal symbiont density and suppression of energy metabolism in the scleractinian coral Pocillopora damicornis. Marine Pollution Bulletin, 191. 10.1016/j.marpolbul.2023.114897

